# MethylSPWNet and MethylCapsNet: Biologically Motivated Organization of DNAm Neural Network, Inspired by Capsule Networks

**DOI:** 10.1101/2020.08.14.251306

**Authors:** Joshua J. Levy, Youdinghuan Chen, Nasim Azizgolshani, Curtis L. Petersen, Alexander J. Titus, Erika L. Moen, Louis J. Vaickus, Lucas A. Salas, Brock C. Christensen

**Affiliations:** Program in Quantitative Biomedical Sciences, Geisel School of Medicine at Dartmouth, Lebanon, NH 03756; Department of Epidemiology, Geisel School of Medicine at Dartmouth, Lebanon, NH 03756; Emerging Diagnostic and Investigative Technologies, Department of Pathology, Dartmouth Hitchcock Medical Center, Lebanon, NH 03756; The Dartmouth Institute for Health Policy and Clinical Practice, Lebanon, NH, 03756; Department of Life Sciences, University of New Hampshire, Manchester, NH 03101; Department of Biomedical Data Science, Geisel School of Medicine at Dartmouth, Lebanon, NH 03756; Department of Molecular and Systems Biology, Geisel School of Medicine at Dartmouth, Lebanon, NH 03756; Department of Community and Family Medicine, Geisel School of Medicine at Dartmouth, Lebanon, NH 03756

**Keywords:** Deep Learning, DNA Methylation, Central Nervous System Tumors, Capsules, Prior Biological Knowledge, Neural Networks, WGCNA

## Abstract

DNA methylation (DNAm) alterations have been heavily implicated in carcinogenesis and the pathophysiology of diseases through upstream regulation of gene expression. DNAm deep-learning approaches are able to capture features associated with aging, cell type, and disease progression, but lack incorporation of prior biological knowledge. Here, we present modular, user-friendly deep learning methodology and software, *MethylCapsNet* and *MethylSPWNet*, that group CpGs into biologically relevant capsules – such as gene promoter context, CpG island relationship, or user-defined groupings – and relate them to diagnostic and prognostic outcomes. We demonstrate these models’ utility on 3,897 individuals in the classification of central nervous system (CNS) tumors. *MethylCapsNet* and *MethylSPWNet* provide an opportunity to increase DNAm deep learning analyses’ interpretability by enabling a flexible organization of DNAm data into biologically relevant capsules.

## Introduction

DNA methylation (DNAm) is a key epigenetic regulator of gene expression in health and disease states, processes of aging and cellular differentiation/stemness, and response to environmental exposures [1–3]. DNAm of cytosine in the context of cytosine-guanine dinucleotides (CpG) sites can be measured with standardized genome-scale oligonucleotide bead-arrays at hundreds of thousands of sites [4,5]. Though a CpG is either unmethylated or methylated, fluorescence signal intensities from array measures of bulk biospecimen DNA are used to derive a beta value measure that approximates the proportion of methylated DNA copies. Gene promoter CpG island methylation is associated with repression of transcription, whereas unmethylated CpG islands are permissive to gene transcription. Alterations to DNAm have a well-established role in carcinogenesis and tumor progression, including inactivation of tumor suppressor genes, aberrant oncogene expression, and loss of repression of repetitive element sequences that contribute to genomic instability [6,7].

The World Health Organization Central Nervous System (CNS) tumor classification includes over 38 tumor types defined by histopathological features [8]. Most of the 38 can be grouped into the broader glioma, ependymoma, and embryonal tumor types. Within those three categorizations, over 80 further delineations are specified by molecular subtyping. DNAm alterations have been heavily implicated in the development and prognosis of CNS tumors. For instance, epigenetic silencing of *MGMT* is associated with an improved response to chemotherapy in glioblastoma patients through the deactivation of crucial DNA repair mechanisms [9]. *IDH* mutations are associated with improved survival in glioma patients through subsequent global hypermethylation of CpG island promoters, known as induction of the CpG Island Methylator Phenotype (CIMP) [10–13]. Other examples include hypermethylation of Wnt and Shh pathways in medulloblastoma patients [14]. The success of differential methylation analyses in characterizing CNS tumors has recently led to the development of DNAm classifiers of brain tumors as companion diagnostic tools to understand and correctly diagnose challenging histologic cases and for the selection of targeted therapies [8].

While the development of this methylation-based brain-tumor machine learning classifier has been heralded as an improvement, existing diagnostic framework clinically applicable classifiers use only a small subset of measured CpGs (e.g., 10,000) [15]. Incorporating additional CpG predictors may allow for the resolution of tumor classes otherwise not identified and help understand relationships with outcomes [16]. This problem may be better approached using machine learning analyses by merit of their prohibitive dimensionality. Deep learning algorithms are a subclass of machine learning approaches that are based on the use of artificial neural networks (ANN) [17–19]. Multi-layer perceptrons (MLP) represent a subclass of neural networks that treat the input data as a one-dimensional vector and then pass the information from one set of nodes to a subsequent set of nodes through fully connected layers of weights/parameters. The information at the subsequent layer of nodes is transformed using non-linear transforms/activations/link functions. These types of analyses are common for deep learning analyses of DNAm data, where the input data is a list of beta values for each subject [20].

DNAm deep learning frameworks, e.g., MethylNet, can accurately characterize tissue, disease states and infer subject age and cell type proportions through unsupervised embedding, generation, classification, and regression tasks [20–24]. They also attempt to ascribe important methylated loci using model interpretability frameworks such as SHAP [25] or LIME [26]. While the inclusion of more CpGs presents an opportunity to expand the space of biologically testable hypotheses [20], statistical challenges (e.g., multicollinearity) with interpretations and generation of associations with pathways remain understudied [27].

Multicollinearity, the unusually high correlation between features, can be addressed with careful feature-selection or grouping [1,28]. Feature selection methods and statistical learning methods such as sparse Group LASSO and network regularization have identified important CpGs in highly complex data [29–33]. More recent work has called for a greater understanding of DNAm-DNAm interactions’ implications through the incorporation of Gaussian Graphical Models, Canonical Correlation Analysis, and module discovery through weighted gene co-methylation networks [34–50]. There is growing support for the use of novel deep learning methods to aggregate, group, and select CpGs by their local context (e.g., genes) to connect and interpret the data with clinical outcomes [51–53]. Incorporation of prior biological knowledge improves the transparency and interpretability of the modeling approach and reduces noise while increasing signal by meaningfully pruning redundant relationships between predictors [54].

Capsule networks have served as inspiration for methods that group CpGs to harness their statistical interactions and relate predictors’ groupings to clinical and biological outcomes [27]. Capsule networks explicitly model the relationships between constituent parts/groups of predictors, or capsules, through parameterizing pose matrices (unitary transformations) and then hierarchically associate each of these parts independently to higher-order targets of interest. While capsule networks are primarily featured in the computer vision domain, evolving methods within different biomedical specialties often utilize grouped organization of predictors in the neural network design [55].

Here we provide a deep learning framework for methylation data that draws inspiration from capsule networks. We investigated the organization of CpG features into DNAm capsules, which represent local contexts that can be related to one another. *MethylSPWNet* and *MethylCapsNet* organize sets of CpGs into a series of capsules defined by higher-order genomic contexts and performs classification tasks (Figure 1A, B). To bring additional interpretability to existing deep learning approaches while capturing hierarchical association networks, we propose and explore *MethylSPWNet* (Sparse Pathway Network) and *MethylCapsNet* as deep learning analogs of these enrichment tests. We provide recommendations for developing these capsule and network-deriving models and provide open-source software for training these models. The *MethylCapsNet* framework proposes to expand the broad utility of these tools by allowing end-users to construct their unique capsules that represent an array of biologically plausible contexts that further explain their target of interest.

**Figure 1:**
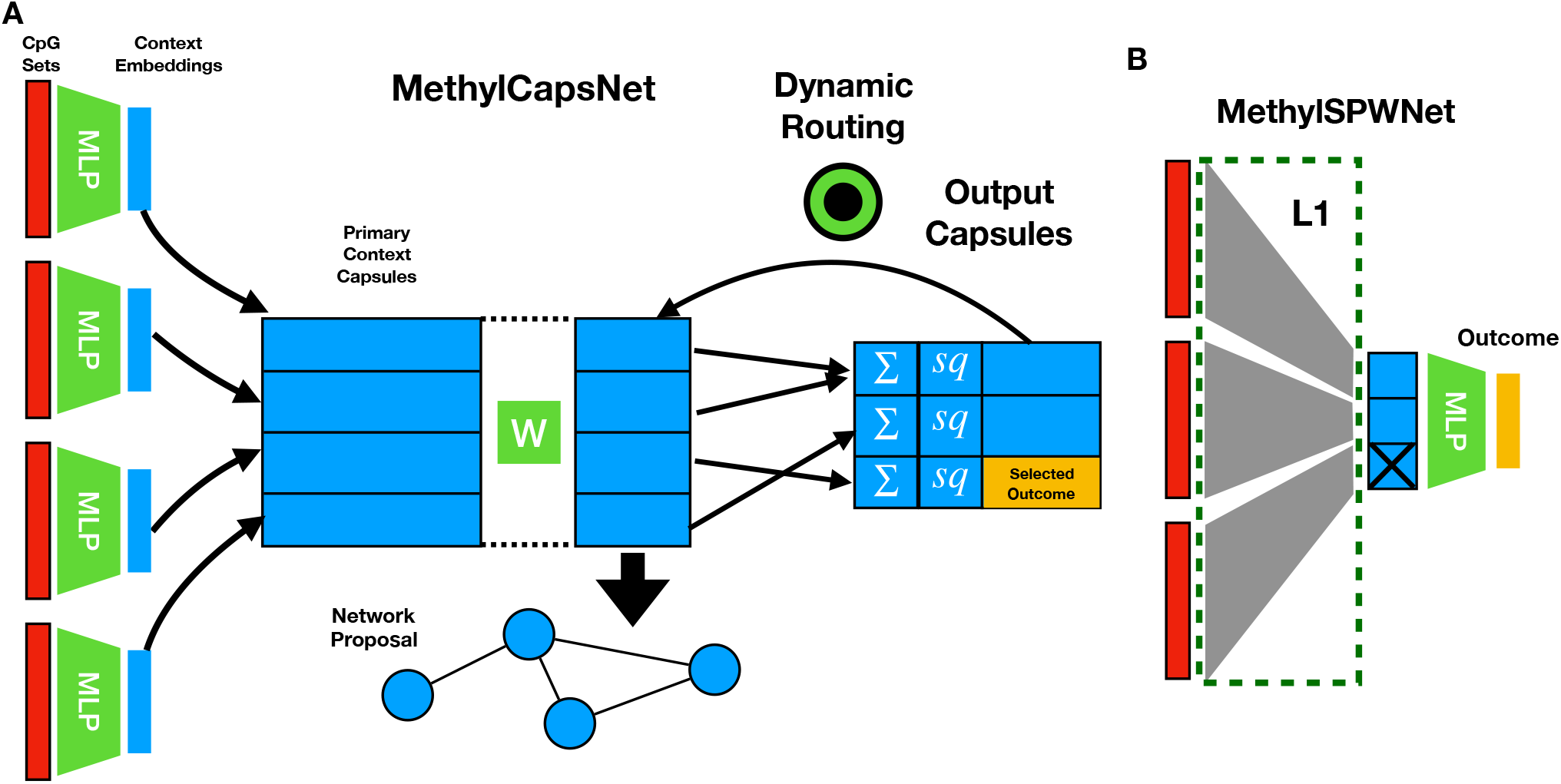
Description of Modeling Approaches: A) *MethylCapsNet* separate MLPs are utilized to form capsule level embeddings from beta values of CpG groupings, which are then associated with outcome targets of interest through dynamic routing of information; these embeddings may be studied to propose networks based on co-methylation on the individual level; B) *MethylSPWNet* aggregates the beta values of groups of CpGs through one locally connected layer; nodes of the resulting layer represent biologically meaningful units that are passed through an MLP for final prediction; group L1 penalties are applied to prune genes/capsules potentially unrelated to the outcome; red colors indicate beta values for CpGs, which serve as input to the algorithm

## Results

To illustrate the potential utility of capsule-inspired neural network approaches, we revisited the Capper et al. [8] dataset used to train a model that differentiates CNS tumors [56]. CNS tumor histology is largely characterized by the presence or absence of morphologically distinct cells of origin, including neuronal, astrocytic, microglial, oligodendrocytic, and Schwann cells. We aimed to predict the 38 histological subtypes of CNS tumors (39 classes, including controls) as a test case for the capsule-inspired neural network approaches. While distinct cell types may characterize these histological subtypes, it was not our aim to classify these cell types through this modeling approach, as methods for brain cell type estimation using DNAm data are still under development. We compare the *MethylCapsNet* and *MethylSPWNet* frameworks for capsule organization with the existing *MethylNet* framework (which does not account for Capsule organized information [20]) and a Random Forest model fit on 10k important CpGs derived using a previously established method (*Random Forest 10k*). Additionally, we provide a Random Forest model on the capsule-organized information extracted from *MethylSPWNet* (*Random Forest Capsules*). Additional details of modeling approaches, fitting procedures, and capsule selection are in the Methods section.

### Capsule Generation for CNS Tumor Prediction

Capsules may be supplied to the neural network approaches in the form of annotations and/or gene sets from MSigDB and GSEA: 1) genes, 2) sites upstream/downstream of the gene, 3) the following Illumina methylation array annotations – UCSC_RefGene_Name, UCSC_RefGene_Accession, UCSC_RefGene_Group, UCSC_CpG_Islands_Name, Relation_to_UCSC_CpG_Island, Phantom, DMR, Enhancer, HMM_Island, Regulatory_Feature_Name, Regulatory_Feature_Group, and DHS – and 4) the following GSEA gene sets: C5.BP, C6, C1, H, C3.MIR, C2.CGP, C4.CM, C5.CC, C3.TFT, C5.MF, C7, C2.CP, C4.CGN. Importantly, users can also input custom capsules into the pipeline through a dictionary that maps CpG to a context name of choice. Finally, capsule generation has been integrated with BedTools [57] (*genomic_binned* selection), which can break up the entire hg19 genome into overlapping windows of fixed width. CpGs in these windows will belong to these capsules. We utilized gene capsules for the primary classification study, though alternative methods for capsule formation are explored in section “Exploration of Alternative Capsule Formations and Cancer Subtypes.”

### Classification Study

We trained each of the modeling approaches to differentiate 38 histological subtypes of CNS tumors and compared their classification performance via a 1000-iteration non-parametric bootstrap of F1-scores over the test set, which balances sensitivity and specificity and reduces the bias in output. Our results indicate that *MethylNet, MethylSPWNet*, and *MethylCapsNet* can achieve very similar high performance on a common data set (Table 1). The neural network approaches achieved marginally better performance than the Random Forest approaches. A breakdown of classification scores for the capsule-inspired models has been included in the supplementary material (Supplementary Table 1). Since all three neural network approaches offer similar performance on classifying brain tumors, we next sought to uncover overlap or complementary insights provided by each modeling approach based on their data organization. The high predictive accuracy of both capsule approaches provided grounds for exploring the factors related to its decision-making process for increased transparency and validation of our approach.

**Table 1:** Classification Results for Random Forest Approaches, MethylNet, MethylSPWNet, and MethylCapsNet*

| *Approach* | *Accuracy±SE* | *Recall±SE* | *Precision±SE* | *F1-Score±SE* |
| --- | --- | --- | --- | --- |
| *Random Forest 10k* | 0.96±0.007 | 0.98±0.012 | 0.93±0.013 | 0.95±0.012 |
| *Random Forest Capsules* | 0.94±0.0087 | 0.97±0.016 | 0.91±0.017 | 0.93±0.016 |
| *MethylNet* | 0.97±0.0061 | 0.99±0.0099 | 0.96±0.011 | 0.97±0.011 |
| *MethylSPWNet* | 0.98±0.0049 | 0.96±0.0088 | 0.96±0.0085 | 0.96±0.0089 |
| *MethylCapsNet* | 0.98±0.0044 | 0.98±0.012 | 0.98±0.01 | 0.98±0.011 |

\*All scores are reported as macro-measures across all subtypes; 95% confidence intervals estimated using 1000-sample non-parametric bootstrap

### Clustering Gene-Level Brain Cancer Embeddings

Until this point, our unit of analysis has been individual CpGs. Summarizing gene-level methylation using median or mean methylation is generally not appropriate. The relationship between methylation state and gene expression can vary depending on the genomic context (e.g., promoter and gene body). However, while training *MethylSPWNet* to predict tumor histological subtype, the model learns to generate gene-level summaries of methylation by updating the weight of each CpG when aggregating beta values of CpGs on the gene-level (See Methods “Description of MethylSPWNet”). This gene-level aggregation can transform a design matrix of samples by CpGs into samples by genes. Gene-level embeddings correlated with outcome (here tumor histological subtype), can then be interrogated for their relationship with known pathways and gene networks. To visualize gene-level embeddings, we generated cluster heatmaps, where rows constitute observations and columns comprise genes, showing plots of the top 2000 variable genes from neural network gene-level embeddings (Supplementary Figures 1-2).

To assess the representation capacity of MethylNet, MethylCapsNet, and MethylSPWNet embeddings, we clustered embeddings with histologic tumor subtype, cell of origin, and histological subtype with the molecular subclass (Supplementary Table 2). Preliminary clustering of the observations demonstrates, for instance, the inability to differentiate *IDH* mutant subtypes of glioma when defined by median methylation versus the neural network parameterization. There is observed concordance between hierarchical clustering in this embedding space and the brain cancer subtypes. This concordance is defined by their molecular subtypes, the original histological subtypes that the model was trained on, or higher-order cells of origin (e.g., mesenchymal, ependymal, neuro-glial origin). *MethylSPWNet* had the highest degree of concordance with histological and molecular subtypes within the gene-level embedding space [58] (V-Measure 0.72±0.0059; Supplementary Table 2). The ability to recapitulate relevant histological subtypes of CNS tumors through the embeddings alone is further corroborated by embedding plots [58]. In these, MethylCapsNet appears to generate the best separation and differentiation of subtypes (Silhouette Score: 0.52±0.0048), followed by MethylNet (Silhouette Score: 0.25±0.01), and MethylSPWNet (Silhouette Score: 0.1±0.0087), estimated using a 1000-sample non-parametric bootstrap. Since these three approaches were trained to recognize histological subtypes, the signal of the cell of cancer origin and molecular subtypes were less well captured.

### Gene-Level and Modularity Enrichment Analyses

Next, we aimed to evaluate the utility of the group-regularized deep learning approach for capsule-organized summaries of DNAm on the gene-level for pathways and gene network analyses. We performed a preliminary analysis of pathway and module detection based on the extraction of hypervariable genes across the neural network embeddings and Louvain clustering of networks of genes based on the pairwise correlation between the genes. Further description of the methods and results is provided in the supplementary material (Supplementary Table 3; Supplementary Figures 3-4).

A description and flow-chart showing an overview of methods for pathways and gene network analysis downstream from *MethylSPWNet* can be found in the Methods section “Description of Potential Downstream Analyses.” We focused our presentation of results on three specific CNS tumor subtypes: glioblastoma (GBM), low-grade glioma (LGG), and medulloblastoma (MB). Gene-level embeddings (gene by sample) and pathways and gene network analysis results (derived from those embeddings) are shown in Figures 2, 3, and 4, respectively. Results on pathways and gene network analyses for these three CNS tumor subtypes (GBM, LGG, and MB) are provided in additional files 1-3. A description of the additional files may be found in the supplementary materials (section “Description of Additional Files”).

**Figure 2:**
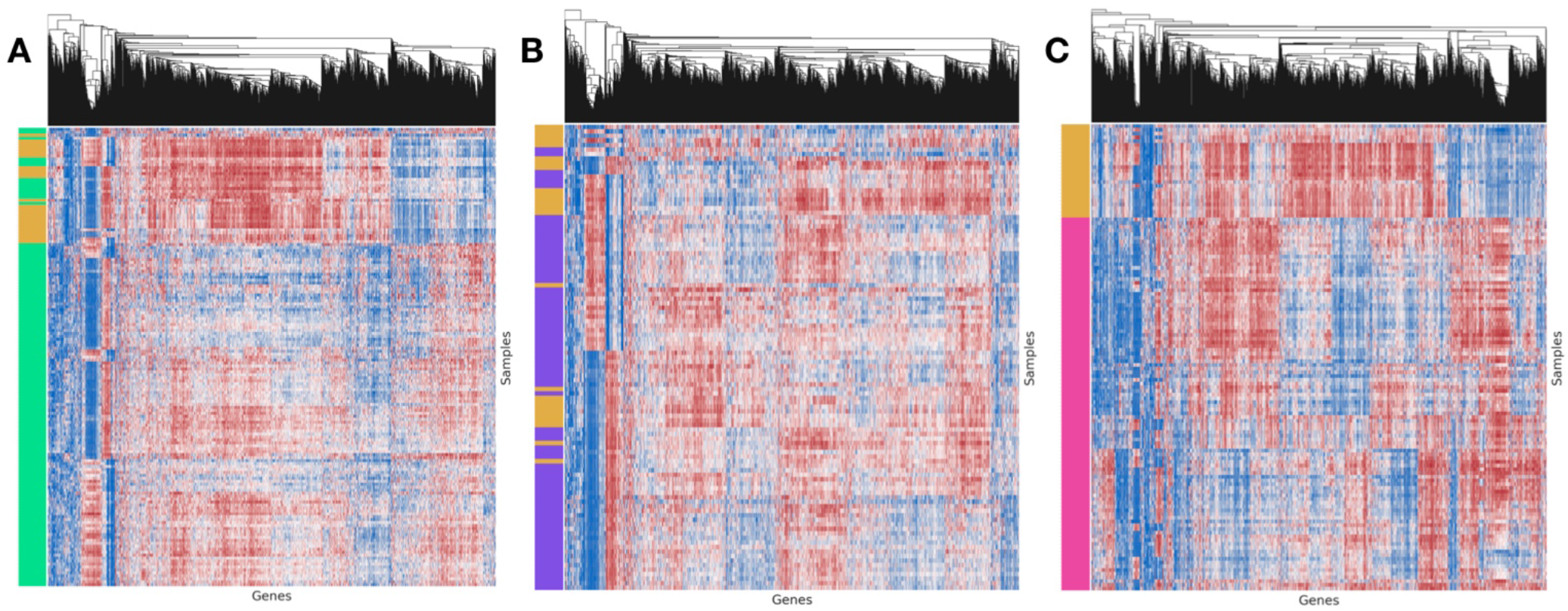
Clustermap of gene-level embeddings for 2000 most variable genes in: A) GBM; B) LGG; C) MB. The left color track indicates the presence of the subtype; yellow indicates the presence of controls; green, purple, and pink indicates the presence of the GBM, LGG, and MB subtypes, respectively; columns have been standardized to highlight trends; in the heatmap, red indicates high *MethylSPWNet* embedding values, while blue indicates low embedding values

**Figure 3:**
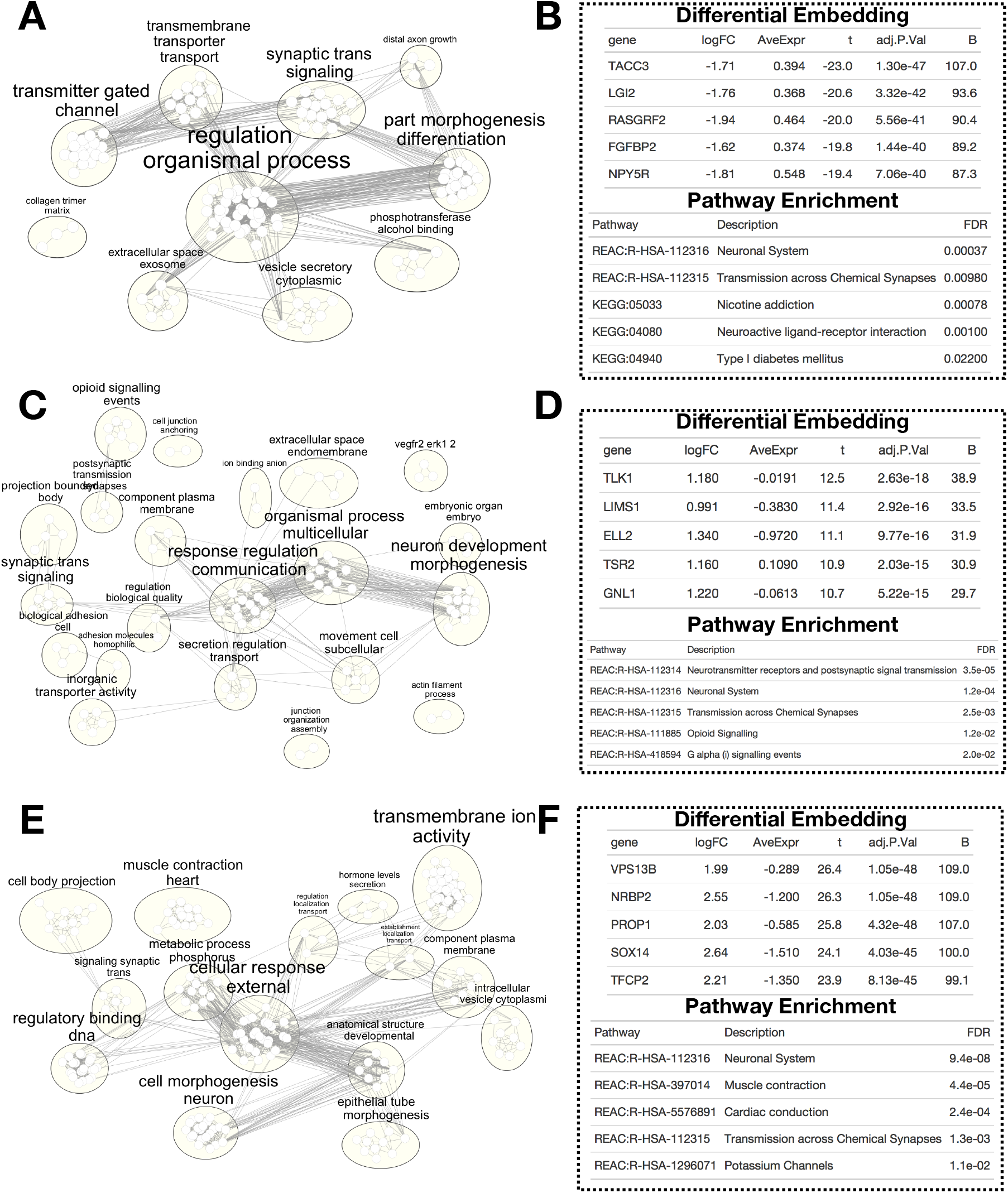
Example Output from Pathway Enrichment Analyses for: A-B) GBM-Control embeddings; C-D) LGG-Control embeddings; E-F) MB-Control embeddings; left-side plots: corresponds to summaries of GO/KEGG/Reactome enrichments using EnrichmentMap in Cytoscape, plotted via RCy3; each small node corresponds to a pathway; node and text size proportional to the number of overlapping genes after differential gene-level embedding analysis; edge between nodes corresponds to shared genes; large ellipses correspond to found clusters using Markov Chain clustering, three words extracted to annotate cluster using a word cloud algorithm; right-side plots: example output of differential embedding (*limma*) and pathway enrichment analyses (g:Profiler) on embeddings; top 5 genes and select pathways listed of the many uncovered in each analysis.

**Figure 4:**
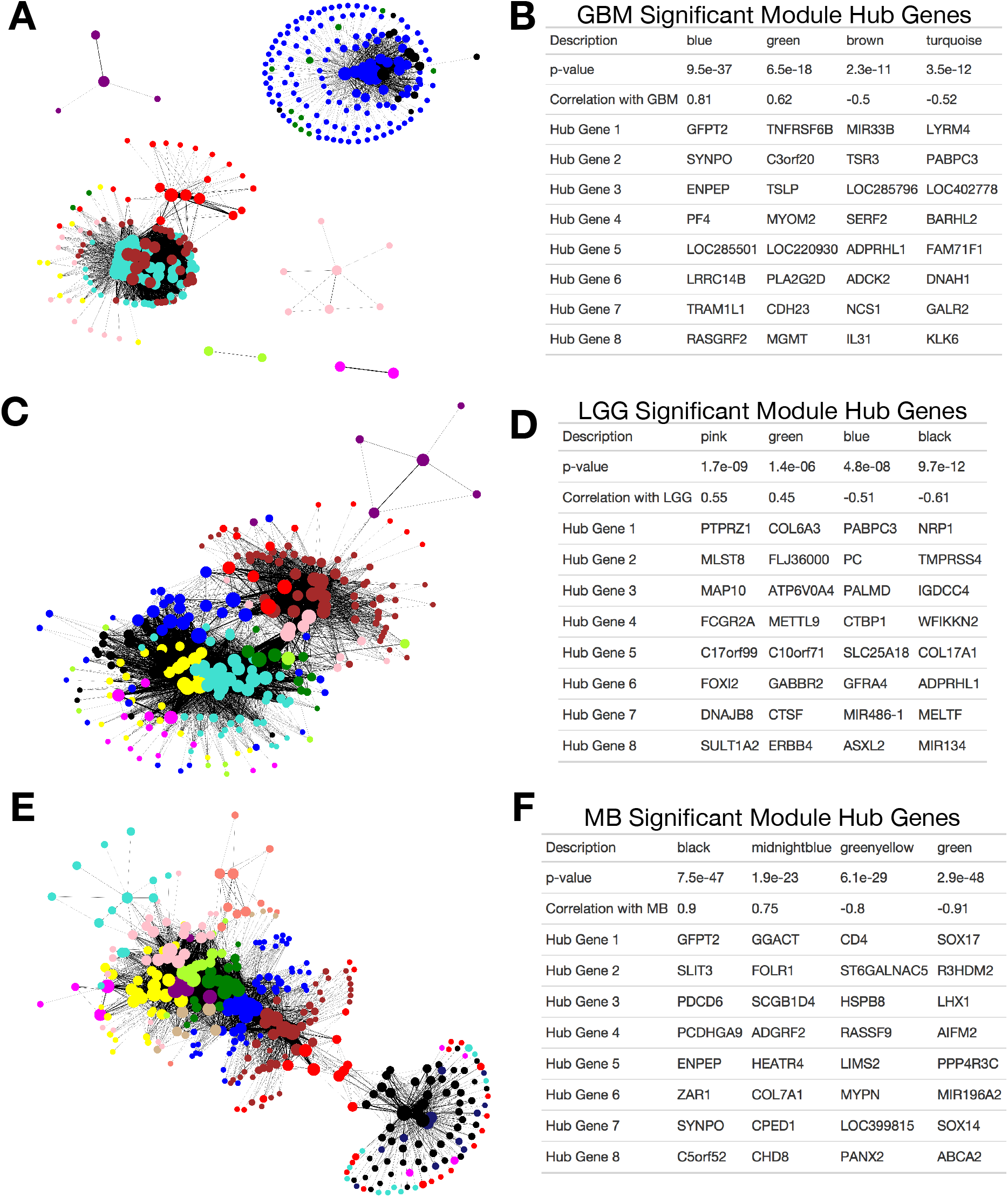
Example Output from WGCNA Analyses for: A-B) GBM-Control embeddings; lack of connectivity between the blue/black modules and the other models suggests lack of functional relationship; C-D) LGG-Control embeddings; E-F) MB-Control embeddings; left-side plots: summarized gene-gene networks using the MAPPER algorithm; nodes indicate genes or dense grouping of genes; colored by discovered module via WGCNA; edges indicate co-methylation; right-side plots: example output from WGCNA analysis; the subset of gene-gene modules; p-value indicates whether significantly associated with tumor subtype; correlation value indicates directionality and strength of this relationship; select hub genes indicate genes with the highest centrality in each subnetwork/module.

### Subtype-Specific Pathways Discovery

Using the gene-level *MethylSPWNet* embedding values, we sought to calculate differentially embedded genes between disease and controls (empirical Bayes) and determine the associations of these genes with some of their correspondent pathways (see Methods “Description of Potential Downstream Analyses”). A visualization of pathway networks and output of gene-subtype associations and pathway enrichment statistics are provided in Figure 3. The empirical Bayes results of the differential embedding analyses for each subtype are provided as Additional Files.

Our analysis of the selected three CNS tumor subtypes (GBM, LGG, and MB) found many top differential genes have been implicated for these subtypes in prior literature. For instance, *TACC* and *FGFBP2* (Figure 3B) are differentially embedded between GBM and controls and have been implicated with a tumorigenic gene fusion event (*TACC*-*FGFR3* [59–61]). *RASGRF2* (Figure 3B) has been linked to congenital GBM [62]. *LGI2* and *NPY5R* (Figure 3B) have both been related to changes in Sox2 expression [63], which promotes cellular plasticity in GBM. An interesting associated pathway for GBM was type 1 diabetes mellitus (Figure 3A-B), an autoimmune disease, of which its spurious association could be related to found associations with several immune evading markers. For LGG, *TLK1* (Figure 3D), a serine-threonine kinase associated with replication, focal adhesion, and cell cycle, has previously been implicated in gliomas [64]. Similarly, low *ELL2* expression (Figure 3D), regulated by microRNA (miRNA)-mediated gene silencing, was reported to be a marker for poorer survival in GBM patients [65]. *GNL1* (Figure 3D) was found in our analysis to be associated with LGG and has been identified as being related to cell proliferation, given its role in the phosphorylation of Rb [66]. We also found associations with the Opioid-signaling pathway and G alpha (i) signaling events [67], and the tyrosine kinase receptor pathway VEGFR (Vascular Endothelial Growth Factor Receptor) and downstream signaling pathway ERK (Figure 3C-D), largely involved with proliferation and angiogenesis [68]. Regarding MB, as examples, we uncovered *NRBP2* (Nuclear Receptor Binding Protein 2; Figure 3F), which had been shown to be downregulated in MB[69], and *SOX14* (Figure 3F), part of the *SOX* family which largely determines cell fate and thus heavily implicated across many CNS tumors[70]. Additionally, pathways such as Muscle Contraction (Figure 3E-F) have been associated with specific molecular subtypes of MB [14,71].

### Grouped-Subtype Pathways Discovery

Additionally, we investigated associations uncovered by grouping together a few select disease subtypes. We would expect, at minimum, differences between these subtypes and healthy controls to be related to pathways that are specialized to those larger histological groupings. First, we compared melanoma-related CNS tumors (MELAN/MELCYT) to controls by performing enrichment analyses of the top 40 differentially embedded genes, as defined by the ranked p-values. As a few examples of potentially enriched gene sets across multiple databases after Bonferroni adjustment, we found potential enrichment for MITF transcription factor targets (*TRRUST*; p=0.06) and neural crest differentiation (*Wikipathways*; p=0.07), BMP signaling (*GO Biological Processes*; p=0.03) and IL23-mediated signaling (*NCI-Cancer*; p=0.05). Of interest from the ependymal tumors (EPN/SUBEPN) was that the top 40 genes had demonstrated overlap with genes related to the spinal cord (*Human Gene Atlas*; p=0.03).

### Derivation of Weighted Gene Co-Embedding Networks

To investigate the gene-level embeddings for each of the 38 brain cancer subtypes (paired to normal controls), we derived disease-specific modules of genes using the Weighted Gene Correlation Network Analysis (WGCNA) R package (see Methods “Description of Potential Downstream Analyses”) [48].

We derived 606 modules of genes across the 38 subtypes (37 networks were derived, one subtype was omitted for low sample count), 297 of which were significantly associated with subtype (all P-values < 0.05). We have included as Additional Files the module membership of each of the genes, module expression across the samples for the three example subtypes (GBM, LGG, MB), hub genes for each module (genes located most centrally in each subnetwork), and statistics that relate each module to the subtype. The connectivity of individual genes from the generated WGCNA modules for GBM, LGG, and MB subtypes are shown in Figure 4. Tables of top hub genes from selected modules strongly associated with each subtype are shown.

Some of the WGCNA modules’ hub genes were found to be correspondent with prior knowledge about their respective subtypes. In GBM, *RASGRF4* (blue module; Figure 4A-B) was previously featured in Figure 3B. The silencing of *MGMT* (green module; Figure 4A-B) plays a significant role in the progression of GBM through inactivation of its DNA repair mechanisms [9]. *MIR33B* (brown module; Figure 4A-B) is also related to GBM progression by regulating cell proliferation, invasion, migration, and MYC signaling [72,73]. Finally, the role of platelet factors (*PF4*; blue module) and CpG island hypermethylation (homeobox gene *BARHL2*; turquoise module) has previously been implicated with GBM [74,75]. Examples of hub genes in LGG include *NRP1* (black module; Figure 4D) [76], *PTPRZ1* (pink module; Figure 4D) [77], and *COL6A3* (green module; Figure 4D). *NRP1* has been shown to be related to poor prognosis in gliomas and signals through microglia/macrophages. *PTPRZ1* has previously been related to malignant growth in GBM. Finally, *COL6A3* is a member of genes serving to form the tumor vasculature. Finally, of interest in MB were *SOX 14* and *SOX17* (green module; Figure 4F) [70], *CD4 (*green-yellow module; Figure 4F*), SLIT3* and *GFPT2* (black module, Figure 4F) and *SYNPO* (blue module, Figure 4F). *CD4* is a gene that codes for the membrane glycoprotein of the CD4 T-cell, where its characterization could be corroborated by the immune infiltration patterns of the stroma for MB. *SOX14* and *SOX17* are pertinent for cell fate lineage. *SLIT3* is a gene characterized by axon guidance and consequently tumor growth, migration, and angiogenesis. Interestingly, *GFPT2* (amino acid metabolism), implicated with higher expression and lower GBM survival [78]). *SYNPO*, central to the black module of MB, were also central to the blue module of GBM.

### Enrichment of Neural Network CpGs for Gene and Island Contexts

As Weighted Gene Correlation Network Analysis (WGCNA) identifies significant associations with known pathways and novel gene-gene co-methylation networks, we sought to investigate the CpG-specific parameters that corresponded to producing the embeddings to understand better why the neural network decided to upweight some CpGs, but not others. To elucidate the genomic contexts that *MethylSPWNet* found to be important, we explored the CpG island context and spatial relationship to the transcriptional start site (TSS) (Methods section “CpG Island/Gene Context Analysis”). CpG islands (CGI) are CpG dense regions. Approximately 60% of gene promoters contain CpG islands[79]. CpG shores immediately flank the CGIs by up to 2kb, shelves flank the shores by an additional 2kb, as regional CpG density decreases. Variables for the spatial relationship to the TSS include TSS1500 and TSS200, within 1500 bp and 200 bp of the TSS, respectively. Additional TSS variables are the 5’UTR immediately downstream of the TSS, the first exon, gene body, and 3’UTR.

Of note, we found that CpGs with positive weights (rank-ordered) were depleted for promoter island regions (defined as having TSS1500/TSS200 annotation and not open sea) (OR=0.69; p=0.04) as compared to sites not included in promoter-island regions (OR=1.45; p=0.04). However, when limiting the set of CpGs to only promoter regions (i.e., TSS1500/TSS200), we noted that positive weight CpGs were enriched for island context (i.e., not in an open sea region) (OR=1.21; p=0.03), while negative weight CpGs were depleted for the CGI context (OR=0.89; p=0.02). Furthermore, both the positive and negative weights of intragenic CpGs were depleted for association with the correspondent methylated promoters (as compared to unmethylated promoters) for their respective genes (positive weight OR=0.54, p<0.01; negative weight OR=0.44, p<0.01). We have included tables for the relationship between CpG weight and independently considered contexts in the Supplementary Materials (Supplementary Table 4; Supplementary Figure 4).

### MethylCapsNet Module Enrichment

From the embedding module discovery analysis and further contextualization of the neural network CpG-weights, we observed that information encoded in *MethylSPWNet* corresponds to key pathways associated with various CNS tumors and important genomic contexts. *MethylCapsNet* offers the ability to infer more granular relationships between capsules on the individual sample level when we can reduce the number of parameters specified. The primary capability and emphasis of the capsule-inspired network approach is to compare capsules to each other and directly relate them to particular outcomes of interest via the dynamic construction of a bipartite network (gene-subtype relationships) as part of the training and prediction process. For the *MethylCapsNet* analysis, we pre-selected a subset of genes previously shown to be implicated in various types of brain cancer (see “Selection of Capsules for *MethylCapsNet* and *MethylSPWNet*” in Methods). As such, we believe it would not be appropriate to test for enrichment of these genes due to the pre-selection procedure that introduces a bias. Instead, we derived modules of genes that the neural network deemed to have a coordinated DNAm response in elucidating particular subtypes. Our modularity analysis projects the estimated bipartite graph (gene-subtype) across samples into a univariate graph (gene-gene), then clusters the graph using Louvain modularity to yield four modules of genes (green, red, blue, yellow) (Figure 5).

Here, we offered an example of the kinds of inferences that can be made from the resultant unipartite network and subsequent clustering. For instance, the yellow module implicates relationships between *WNT3A* and *EGFR*, heavily implicated in Igf and Wnt signaling. The red module features the relationship between *FRZ* and *APC*, both of which are heavily involved in WNT signaling (*APC* forms the complex to inhibit the accumulation of β*-catenin*, while WNT binding to frizzled family receptors may degrade this inhibition and permit cell proliferation [80]). Of the green module, *IDH3G* and *NPR3* (linked to energy metabolism, gene fusion, and chromatin remodeling) were related to both *LRDD* (pro-apoptotic *MAPK* pathway) and *WIF1* (both previously implicated WNT signaling suppressors) from the yellow module. *KIAA1549*, related to astrocytomas and fused to *BRAF* for its progression to oncogenesis, was implicated with *WNT1* in the blue module [81]. Insulin-like Growth Factor Binding Protein 2 (*IGFB2*; glioma oncogene) of the red module appeared to be negatively correlated with many of the genes across the yellow and green modules[82]. *TP53* (which lacked consequential methylation patterns) and *MYC* [83], *MGMT*, and *TERT* share relationships with each other but not with the other modules, perhaps highlighting how ubiquitous these somatic alterations are for oncogenesis.

**Figure 5:**
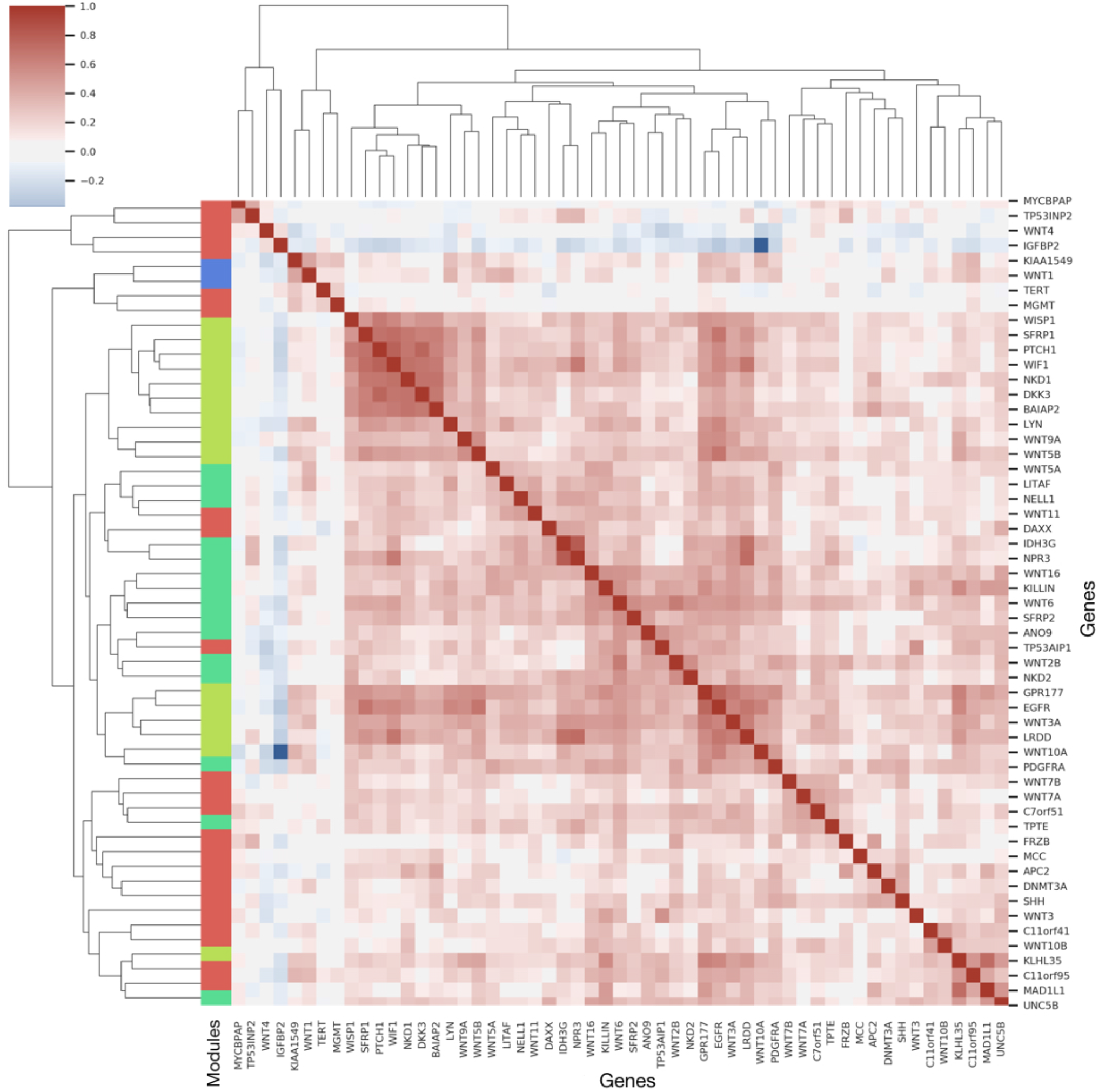
Clustermap of weighted bipartite network projection of select gene capsules’ relationship to various CNS tumors using the *MethylCapsNet* modeling approach; rows and columns are genes, row colors denote modules (green, red, blue, yellow) of genes found using Louvain modularity; values in matrix denote Pearson’s correlation between genes

Despite having highlighted potentially interesting relationships, we acknowledge that there will be a future opportunity to increase the search space of possible relationships between genes and their coordinated response for bringing about subtypes of brain cancer. In the Supplementary Material, we provide the routing matrix that was used to align each gene to a correspondent CNS subtype, which is the predicted gene-subtype bipartite network averaged across individuals (Supplementary Figure 5). We have also provided a locally deployable web application that allows the user to interrogate their uncovered capsules and form networks on the individual level and aggregated across patient subgroups or disease subtypes using gene-capsule-specific embeddings (Supplementary Figure 6). For instance, in our supplemental material, we demonstrate how the *MethylCaps* web application can be used to derive individual networks for LGG and GBM, identifying *ANO9* and *EGFR*, respectively, as implicated in these conditions (Supplementary Figure 7-8). These genes and their associated pathways have been heavily implicated in tumorigenesis and gliomas [84,85].

### Exploration of Alternative Capsule Formations and Cancer Subtypes

In this section, we briefly present results from a few of many alternative means of forming capsules. Particularly, we consider the following scenarios for CNS tumor classification: 1) *MethylCapsNet* is fit when a) only half the genes are retained from the original list (selected randomly) and b) none of the genes are retained and instead randomly sampled from all genes and 2) *MethylCapsNet* is fit using binned genomics regions, utilizing CpGs encapsulated in 1Mb bins. In addition to these analyses, we also explore an integrated breast cancer dataset utilized for PAM50 molecular classification[20,86,87] using a few different capsule configurations: 1) *MethylCapsNet* is fit using a curated list of genes, 2) *MethylCapsNet* is fit using binned genomics regions, utilizing CpGs encapsulated in 700kb bins, and 3) *MethylSPWNet* is fit using capsules organized by CpG island promoters, formed by intersecting CpG island with gene promoter annotations from the Illumina 450k database. A complete set of results can be found in the supplementary material (Supplementary Tables 5-6; Supplementary Figure 9).

## Discussion

Recent reviews and initial explorations discussed the potential utility of capsule-inspired networks to relate biologically-organized capsules to each other and known disease outcomes [27,88–90]. In this work, we set out to perform a preliminary evaluation that shows the feasibility and suitability of DNA methylation capsules for deep learning analyses as a means to organize CpG information to higher-order contexts to improve prediction and transparency while uncovering instances of coordinated gene-level methylation patterns. In our analyses, we compared several state-of-the-art predictive modeling methods for DNA methylation classification of brain tumors. We demonstrated that capsule-based deep learning approaches could achieve performance on par with existing deep learning models and prove better than existing traditional machine learning frameworks for analyzing DNA methylation data. Our work demonstrated the potential for new insights compared to other existing methylation-based tumor classification schemes currently used, which are often based on a small subset of CpGs, and lack built-in interpretation of the loci selected[8,91]. We demonstrated the efficacy of increasing the use of the available CpGs on the Illumina 450k Array, ultimately using 200,000 loci before sub-setting by context.

DNA methylation capsules focused on the gene level can disentangle important CpGs that might otherwise be down-weighted in a feature-by-feature deep learning unsupervised or supervised learning approach. These CpGs demonstrated substantial overlap with genes known to be related to tumorigenesis in the brain, such as *NOTCH1, PTEN*, and *GNAS*. This is consistent with previous studies that demonstrated mutations common in brain tumors, such as *IDH1*, are correlated with disruptions in methylation.[10,92–97]

The context-specific CpG weight enrichment analyses suggest that within promoter regions, island context is important for differentiating different CNS tumor subtypes, but taken as a whole, regions outside of CpG promoter islands are important for capturing this heterogeneity. Furthermore, outside of the promoter context (supposedly regions that better capture tumor heterogeneity), the ability of intragenic CpGs to distinguish tumor subtypes is still dependent on the promoter methylation status of the respective gene. Clustering of CpGs with the highest weights at CpG islands, shores, gene bodies, and transcription start sites will help us understand where the most diagnostically relevant sites are in the genome, but demands additional investigation.

We also presented a few examples of the potential downstream applications of capsule-based approaches. In particular, our framework demonstrated the ability to relate derived gene-level measures of *MethylSPWNet* to known disease pathways via differential methylation analysis of the gene-level embeddings and gene-gene co-methylation networks via WGCNA. Additionally, we provided a preliminary interpretation of bipartite (gene-subtype) and unipartite (gene-gene) networks, which can be derived by *MethylCapsNet* web framework. Finally, we explored alternative means from which to form capsules. We expected the curation of genes to lead to more accurate models. Contrary to our initial hypothesis, for CNS classification, random capsules’ selection appeared to still produce a highly accurate model. These results suggest either the potential to uncover novel associations between genes and subtype or that these genes may be co-methylated with other genes that have well-established relationships. The binned genomics, fit using *MethylCapsNet* for classification tasks in brain and breast, were similar to the leading methods. The island promoter capsules slightly underperformed, suggesting that these capsules’ selection alone does not contain enough information to distinguish PAM50 molecular subtypes.

There are a few limitations to this work, presenting room for future improvements in the analytical method. First, the included CpG loci selection is biased on limited sites available from the Illumina Array platform. At present, the utilization of an Illumina methylation array platform is more tractable due to lower technology costs and expertise required [98] than Whole Genome Bisulfite Sequencing (WGBS) that requires substantial sequencing depth on the order of 100x for comparable precision [99].

Second, a reduced set of genes were fit using the capsule-inspired network. It remains challenging to run MethylCapsNet at scale due to the heavy computational demand and the large number of free parameters in its current formulation. Since *MethylCapsNet* can only analyze approximately one thousand capsules at a time, the capsule selection step is critical to the method’s successful application. This parameter space should be reduced by finding some marriage between the scalability enabled by *MethylSPWNet* and perhaps greater transparency offered by *MethylCapsNet*. Presently, we advise end-users to utilize *MethylSPWNet* when the number of contexts under evaluation is large (≥1000 capsules), or if the number of CpGs per gene is small, and to utilize *MethylCapsNet* when the number of contexts under consideration is smaller (<1000 capsules). Uncurated gene sets can be analyzed using *MethylSPWNet*, while curated gene sets are best suited to *MethylCapsNet*, e.g., regions of the genome fragmented by consistent windows or larger DNAm CpG modules that have been uncovered through methods such as WGCNA. In addition, the adoption of capsule-inspired approaches that explicitly form networks via their routing mechanisms presents a future area of research [89]. It is also assumed that *MethylCapsNet* capsules that are more closely embedded are interacting, but it is not entirely clear the nature of these interactions without incorporating gene expression data and methylated Quantitative Trait Loci analyses.

While inspired by capsule networks, we also emphasize that these methods are not analogous to capsule networks featured in computer vision tasks. While the new capsule-based approaches were as accurate as fully connected approaches, this was done so under the constraint of sparse connections, where such specification points to the validity of imposing these constraints. Given the nature of the problem (classification amongst dozens of histological subtypes), intermediate embeddings may reflect a more linearly separable subspace to subtypes of origin. Such a subspace may require additional exploration/penalization to avoid potential biases pertaining to the minimal redundant set of predictors to produce a subspace optimal for prediction. The application of such methods does not preclude the potential for selection of genes due to technical reasons such as noise, batch effects, and weight initializations, which are common to many domains of application of neural networks. We attempted to account for such biases through preprocessing methods on the data such as functional normalization and note that strict interpretation of threshold cutoffs for methods devised for differential gene expression may not be applicable. Thus, relaxation of the scope of features input into a pathway and other enrichment analyses may potentially reduce bias so as long as limitations are appropriately stated. Additionally, differences in tissue preparation (frozen, permanent) were not accounted for, and however, given the high concordance between these preparation methods, we felt that such adjustment was not necessary [100].

While DNAm deep learning methods with built-in interpretability do not yet exist, we hope these methods, though constrained by potential limitations in design choice, may spur further research into more interpretable capsule methods. Here, there is also an opportunity to further apply concepts from topological data analysis (TDA), such as Mapper[101–105], to distill the key functional relationships from high-dimensional, complex data.

Existing classification frameworks currently used in the clinical setting for aiding brain cancer diagnosis only utilize a small subset of the total possible set of CpGs that can be measured. Current modeling approaches can be difficult to trust or use to study new network biology until they can consider a larger, more complete set of predictors. However, it is also important to note that doing so would introduce additional noise into the modeling approach, but the incorporation of prior biological knowledge can potentially help reduce noise while improving the detection of biologically relevant signals. We note that underperformance could suggest selecting capsules that may not be optimally aligned to the target task/dataset. By demonstrating that the organization of CpGs into their respective genomic contexts, we present further opportunity to reduce the feature space and disentangle correlation and collinearity between CpG sites to create a new class of transparent, clinically tractable models. For instance, future classifiers should include brain cell-type classification using DNAm data and incorporate it as covariates in the prediction model, yet brain cell-type differentially methylated regions for deconvolution by DNAm patterns is not well-established. The opportunity space of epigenetics research questions is ample and poised to grow substantially as the field moves to expand reference-based approaches to cell type deconvolution, include tandem assessments of other cytosine modifications (hydroxymethylcytosine), and apply DNA methylation age clocks to questions of biological aging. Despite having demonstrated the promising downstream analyses that users may readily adopt through our framework, we acknowledge that there is ample opportunity to develop related methods and their use cases further.

## Conclusions

Here, we have demonstrated the feasibility and utility of DNAm based capsules for performing disease classification and potentially determining dysregulated genes for these diseases. We found that DNA methylation capsule methods can predict brain cancer subtypes with high accuracy and present convenient means for organizing data over traditional techniques for studying DNA methylation data. As such, we advocate for the organization of well-defined DNAm capsules as a means to improve the accuracy, transparency, and broad applicability of DNAm deep learning models. Future deep learning prognostic models that reimagine the formation and incorporation of DNA methylation capsules, paired with cell type inference, gene expression, and/or corroborating chromatin capture, may serve as grounds for the derivation of unknown heterogeneity.

## Methods

### Overview of Framework

The *MethylCapsNet* methodology presents an extension of the *MethylNet* framework [20] and is implemented as a command-line interface that allows the user to group CpGs into capsules and then dynamically route the capsules to make a prediction and interpret the results. While this approach draws inspiration from capsule networks featured in computer vision tasks, *MethylCapsNet* is not explicitly a capsule network, as defined in previous works in this domain.

*MethylCapsNet* utilizes separate MLPs for every set of CpGs (one set per context) to derive context-specific embeddings (separate context embeddings per each individual), and dynamic routing processes force information from child capsules into disease/categorical outcomes (Figure 1A). The information is hierarchical because each child capsule may only align with one parent capsule. Once the capsule-inspired network is fit, graph structures that describe the relationships between each individual’s contexts can be derived by thresholding the correlation between pairwise n-dimensional context embeddings. Highly conserved biological networks can be derived by thresholding the number of individuals that share the same edge between the contexts. The simplest genomic context considered are genes that the CpGs annotate to. Other capsules can be defined by, for instance, genomic region or pathway/biological process annotation. *MethylSPWNet* is a specialized neural network architecture that routes beta values from the CpGs in each context into a single node representing the context (Figure 1B). Each CpG is given a weight based on the importance of its contribution, both on the gene level and towards the classification task as a whole. This information passes through additional neural network layers that dynamically relate latent sets of predictors to outcomes of interest, whether they be prognostic or diagnostic [53,106]. Much like Group LASSO approaches, group L1-penalization can be utilized on the CpG weights routed to each gene to select relevant genes of interest.

The software implementation (Figure 6) is comprised of modules pertaining to prediction and interpretation tasks, which take into account the relationships and embeddings derived through the training process.

**Figure 6:**
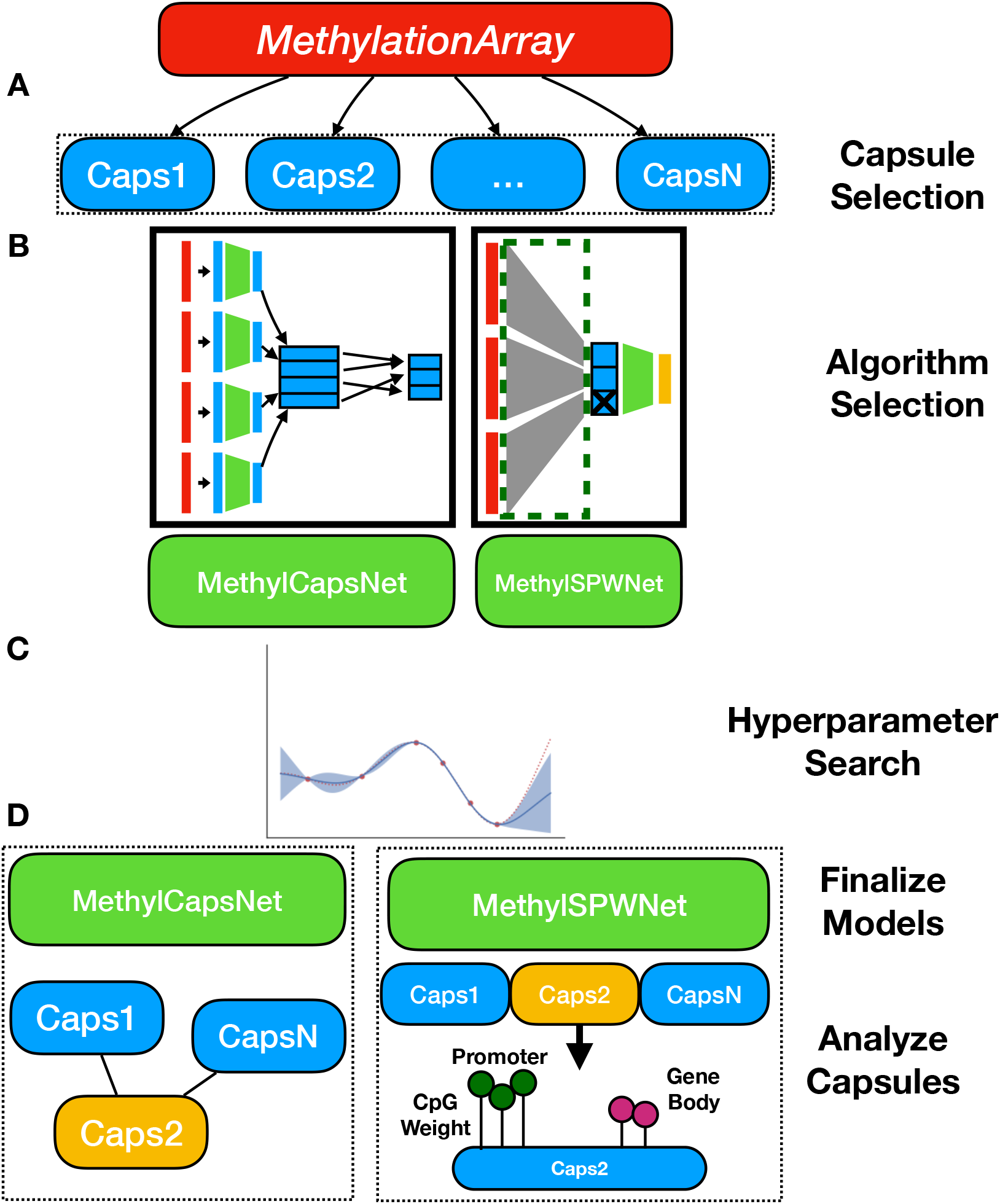
Description of Framework: a) User selects capsules to group sets of CpGs from their own supplied list or a prespecified annotation set; b) User selects modeling approach, *MethylCapsNet, MethylSPWNet, Group LASSO*; c) Hyperparameter search conducted to reveal ideal model specification; d) Final models are fit, and capsules are interrogated for relationships with one another, relationships to the outcome, and highly weighted CpGs; yellow indicates an important capsule; *MethylCapsNet* relates the Capsule to each other per individual; *MethylSPWNet* locates important CpGs within a capsule and contextualizes its location; input data into the algorithm is colored red

### Data Preparation

DNAm data from CNS tumors (n=3,897) were accessed from the GEO archive (GSE109381), preprocessed using *PyMethylProcess* [21], and divided into 70%/10%/20% training, validation, and testing sets (*MethylationArray* objects) via *PyMethylProcess*. The 200,000 most variable CpG loci across the training samples were retained for analysis. Sets of CpGs were tracked to genes, which were then selected to form capsules. The original set of 200,000 CpGs was used as features for the *MethylNet* approach, the complete set of intersecting gene capsules with more than five associated CpGs were used for the *MethylSPWNet* (n=10,341; 139,028 CpGs), and Group LASSO approaches, and a reduced set of capsules (n=55), were utilized for *MethylCapsNet* after manual curation and a hyperparameter search (see “Selection of Capsules for *MethylCapsNet* and *MethylSPWNet*”).

### Description of Potential Downstream Analyses

After fitting a *MethylSPWNet* model (and *MethylCapsNet*), the user may further interrogate the gene-level embeddings depending on the research question being addressed. The user may explore how each gene relates to each outcome or how they relate to one another, the details of which have been included in an informative flow diagram (Figure 7) (separate text describing information that can be extracted after fitting *MethylCapsNet* can be found in the section “Description of Capsule-Inspired Neural Network”).

**Figure 7:**
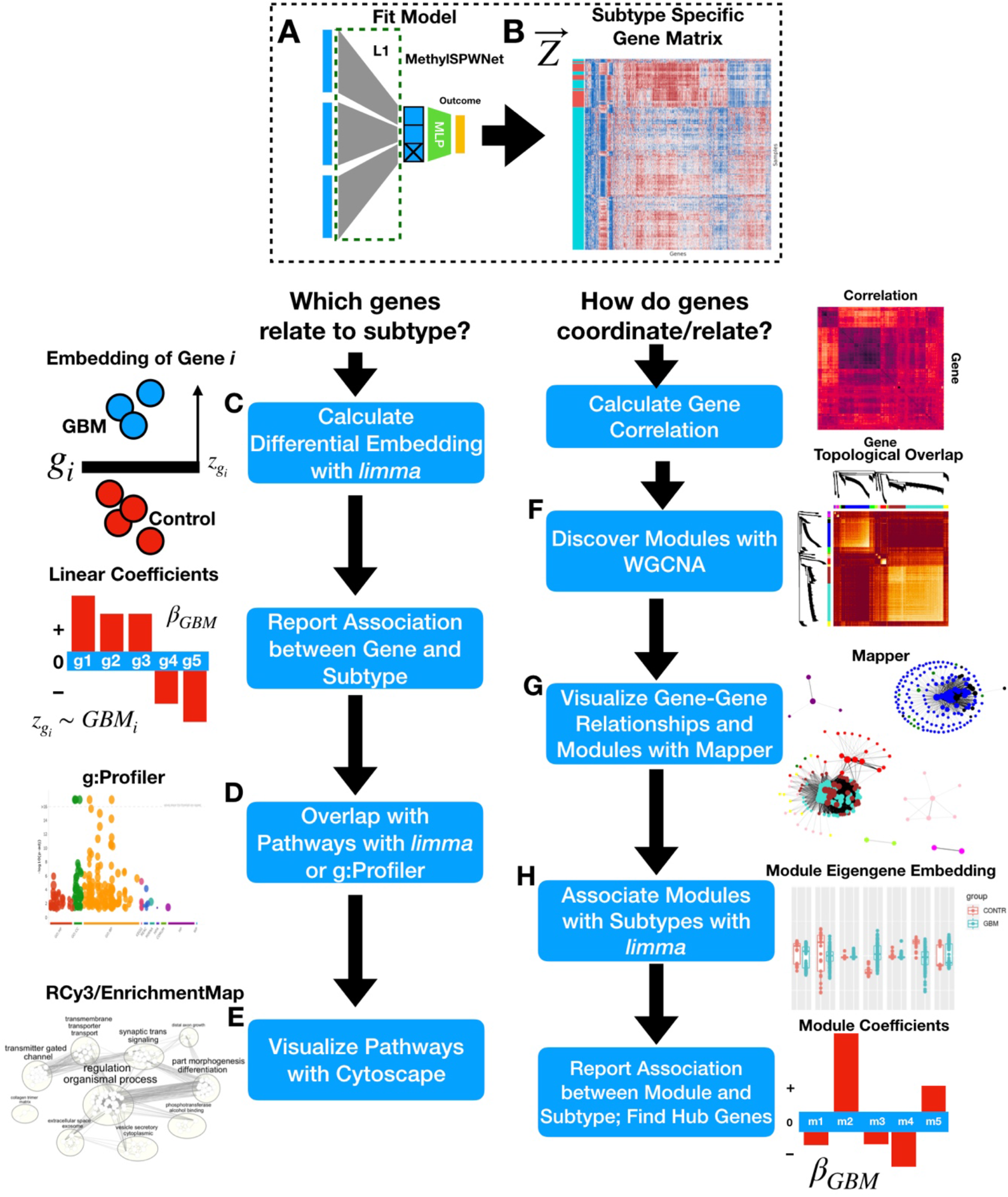
Flow Diagram for Possible Downstream Applications of *MethylSPWNet*: A) User fits *MethylSPWNet* model to predict brain cancer subtypes; B) gene embedding matrix (samples by genes; in this example GBM vs Control) is extracted from the model from the test samples; user decides whether they want to **(C-E)** associate gene-level embeddings with disease outcome, or **(F-H)** relate genes to one another to imply functional relationships; C) user opts for pathways analysis; *Empirical Bayes* method is used to identify genes with differential embeddings between GBM and controls using an empirical bayes moderated linear model; D) significant genes may be passed into g:Profiler or *limma*’s internal functions *camera, goana*, and *kegga*, for pathways enrichment over GO, KEGG, Reactome, etc. databases; E) pathways are summarized and visualized using EnrichmentMap, accessed via RCy3; F) user opts for gene correlation analysis (WGCNA), first calculating correlation between genes, then calculating topological overlap over correlation transformed by power, hierarchical clustering is applied to deduce modules; G) gene-gene networks and module membership is visualized using Mapper; H) modules related to GBM are extracted using the *Empirical Bayes* method by comparing projections into first eigengene (samples by modules matrix) for each module for GBM vs controls

Differentially embedded genes (the extent to which gene-level embeddings vary between subtypes versus normal) from gene-level values derived by the neural network embeddings in each of GBM, LGG, and MB, were identified using the *limma* [107] package. This package compares tumor to nontumor control tissue through least-squares regression and empirical Bayes moderated F-tests, yielding FDR-adjusted p-values and log-odds ratios for the degree of differential embedding. We profiled functionally enriched pathways using the g:Profiler package [108] after selection of genes below an FDR-adjusted significance threshold and visualized the results (relating pathways by the number of shared genes, clustering into higher-order pathways via Markov clustering) using EnrichmentMap, as part of the Cytoscape network visualization framework [109,110].

The pairwise correlation between *MethylSPWNet*-derived gene methylation was calculated using Pearson’s correlation coefficient. Weighted adjacency matrices were calculated from the pairwise correlation matrices for each of the subtypes using the power adjacency function, which takes the co-methylation to a power specified separately for each subtype. To further cluster the genes, the weighted adjacency is transformed into a topological overlap matrix (*TOM*), defined by the extent to which two genes share a third common gene. Finally, hierarchical clustering is applied to derive the final modules of genes. Finally, to relate each module with the disease subtype via the aforementioned least squares and empirical Bayes differential analyses methods, we calculated eigengenes (1^st^ principal component) for the genes in each module to further reduce the design matrix (samples by modules) [107]. A large number of genes (on the order of five thousand) for such a summary gene network plot may make plotting the individual genes cumbersome and hard to understand, so we utilized Mapper[101,103,104], a tool from Topological Data Analysis, to further summarize and portray the relationships between the genes in the network summary plot.

### CpG Island/Gene Context Analysis

Using the MethylSPWNet, each CpG is assigned a weight that relates the CpG to its associated gene or genomic context. These CpG weights are learned by the neural network and can be used to rank genes based on their relative importance (rank assigned by maximum absolute CpG weight), an alternative measurement to the modularity analysis. Inspection of the weights of CpGs within each gene can provide insight into sites and contexts that are important for predicting brain cancer subtypes. Further, investigation of weights spatially across the genome may give rise to important patterns and motifs that could warrant future investigation. In the supplementary material, we first considered the contexts mentioned above independently and did not consider the joint impact of context (e.g., did not associate with island-promoter regions, which are generally considered to be regions more causally related to changes in their expression). We then considered sites that were associated with island promoter regions (including shore and shelf context; more causally associated with gene expression) and separately compared the overlap of the CpGs correspondent to top positive and negative weights to the CpGs that were unassociated with this context (open sea and not TSS200/1500). We separately considered CpGs within the promoter regions. Finally, we considered intragenic CpGs and whether or not their corresponding gene’s promoter was methylated or unmethylated (as operationalized by calculating a beta-value methylation cutoff via local minima in the distribution of beta-values. The beta-value distribution, bimodally distributed, reflects the distribution of proportion of methylated alleles across a bulk mixture of cells for individual CpG sites. This distribution across CpG sites is typically estimated per individual(s). Beta values can take on values between 0 and 1, but particularly concentrate closer to 0 or 1 to reflect that a site is either “methylated” or “unmethylated”. The intermediate proportions reflect scenarios from which around half of the cells of the mixture are methylated at that site, which is uncommon, and used to as the threshold to denote whether a site is methylated. Any CpG with a beta value above the threshold was methylated. CpG methylation was averaged across each promoter and subject to the threshold to determine methylation status). We calculated odds-ratios for enrichment/depletion in these contexts using Fisher’s exact tests. Without matching gene expression information, we could not make any causal claims/inferences about how these contexts modify gene expression to bring about these disease states.

### Description of Capsule-Inspired Neural Network

The capsule-inspired network featured in this work operates by first finding representations of the given CpG sets as denoted by the primary capsule formation. The features, CpGs, of the CpG sets are fed into parallel implementations of a multi-layer perceptron, *f*_*j*_, where the output dimensions of each of the neural networks are the same. Thus, the dimensionality of the primary capsules reflects the number of output neurons, a latent representation of each CpG set, times the number of capsules, per individual. The mathematical formulation of this transformation is presented below:

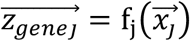

For a single individual, the capsules, represented by row vectors, are stacked to form a capsule matrix:

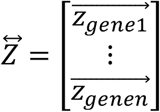

An affine transformation transforms the primary capsules, a set of learnable parameters that seek to rotate, scale, shift the data, and encode information pertaining to the interactions between capsules.

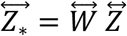

Each primary child capsule’s information is then dynamically routed to parent hidden or output capsules.

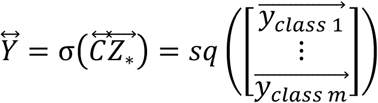

Where:

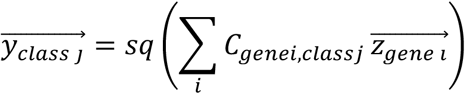

Dynamic routing aims to force the information encoded into each child to align with one parent capsule, thus utilized to calculate 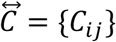, a bipartite network relating the child-parent-capsule. A vector of the same length represents each child as the output of the parent capsule. Analogous to the non-linear transformation of the sum of the information output from the previous layer of neurons for traditional neural networks, for each parent capsule, the child-capsule values are summed, and then a non-linear transform called a squash function, 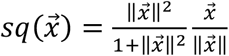, is applied to effectively zero-out, or *squash*, child capsules that do not agree with parent capsules.

Each child’s contributions to a parent are weighted, but two constraints are imposed: First, the weights from each child to its parents must sum to 1. Second, a reward for the alignment of a child to exactly one parent is a dynamic routing by agreement mechanism. An iterative process updates the weights between the child and parent by adding their dot product. The update mechanism for calculating 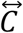 is recapitulated below. After initializing 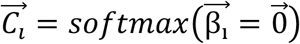 for *r* ∈ {1,2,3, …} iterations:

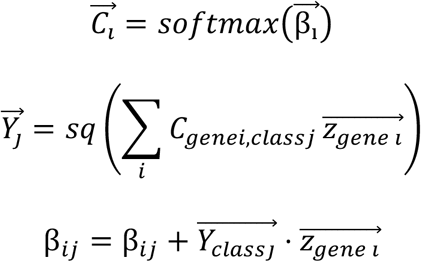

This formula is simplified from its original derivation and utilizes a few notational shortcuts.

Applying this operation for r iterations per batch per training epoch effectively prunes the other connections between the child and its parents as it converges on a single parent from which to send its information. Each output capsule per individual, 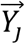, is represented by a vector in some n-dimensional space. The output capsule with the highest L2-norm, 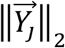, is selected as the predicted class, and a margin loss is applied to penalize the model when it fails to either concretely have a very high (*m*^+^ = 0.9) or very low probability (*m*^−^ = 0.1) of prediction.

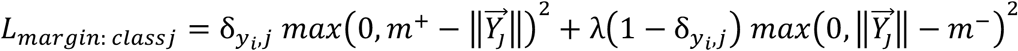

The Kronecker’s delta 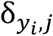 is equal to one when the outcome for individual i is equal to the *j*-th class, thereby activating the left-hand margin loss that penalizes the model if the probability is below *m*^+^ = 0.9. On the right-hand of the equation, 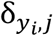 is equal to zero when the outcome for individual i is not equal to the *j*-th class, thereby activating the right-hand margin loss that penalizes the model if the probability is above *m*^−^ = 0.1.

The model is also penalized based on how much the original methylation array could be constructed from the true class’s output capsule via a decoder neural network, 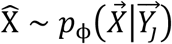:

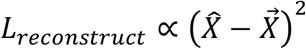

Of the most interest to a biologist may be the primary capsule embeddings per individual, 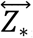, which demonstrate interactions between these biological hypotheses and how the outcome of interest is separable within certain genomic context, and the weights between the primary and output capsules, 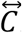, a bipartite graph demonstrates how these genomic regions are related hierarchically and have implications for parent processes. The coordinated response of capsules can also be derived through a bipartite projection of 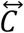 into a unipartite network of capsules. Secondarily, of importance are the concatenation of the primary capsules, which demonstrate overall class separation, and the decoded output. Tweaking the embeddings or L2 norm of the output capsules and decoding can potentially effectively generate methylation data conditionally on outcomes of interest and interpolate between purified states, though this aspect was unexplored due to prohibitive dimensionality.

### Description of MethylSPWNet

*MethylSPWNet* is the deep learning analog of a Group LASSO Regression model. The beta values for the CpGs for each gene, 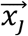, is transformed into a single value, *z*_*genej*_, through the multiplication of a set of gene-specific CpG weight matrices, 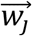. These weights are updated throughout the training process to minimize the divergence between observed and expected outcomes. The magnitude of the weights dictates how much information from each CpG should be considered. The final gene-level summary value is given by:

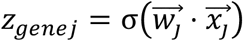

The gene-level summary values are concatenated to form an array of gene-level summaries 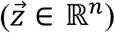:

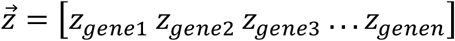

The final prediction for the network can be obtained using the following transformation via an MLP, *f*:

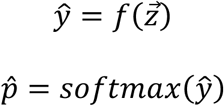

In the classification case, this predicted outcome is compared to the expected outcome via:

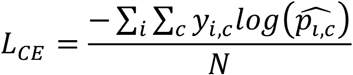

We applied Group L1 regularization to these weights to cause certain genes to drop out, returning genes important for the prediction of the cancer subtypes. The final LASSO penalty is given by:

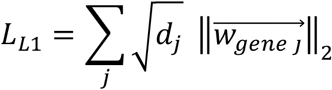

Where *d*_*j*_ is the number of CpGs assigned to that gene. An intermediate layer of the neural network, 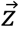, stores gene-level summaries of DNAm information, and 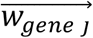 contains the importance of each CpG for a particular gene.

Here, we contrast this summary measure, 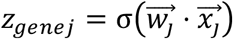 to a more traditional summary measure, such as the median or mean methylation (mean displayed on the right 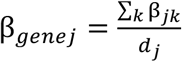. Assuming the mean as our measurement, for simplification, it can be seen that each CpG is given equal weight 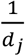, while for *MethylSPWNet*, each CpG is given weight *w*_*jk*_, which is learnable and reflective of the relative contribution of the given methylation beta value to the aggregate measure. A comparison between *z*_*genej*_ and β_*genej*_ can be found in the supplementary materials.

### Hyperparameter Scans

*MethylCapsNet* includes the use of a hyperparameter optimization scheme, accessible through the *methylcaps-hypscan* module. Currently offered by the package is the availability to scan a number of hyperparameters, including the number of training epochs, length of the genomic region, the minimum number of CpGs to constitute a capsule, weighting schemes for reconstruction loss and survival loss, learning rate, in addition to other focused hyperparameters. Additionally, the search of *MethylNet* model architecture, randomized neural network topologies, was replaced by a framework that searches for the ideal number of neurons per neural network layer, conditional upon the choice of the number of layers. There are three search strategies for optimization, including randomized searches and Bayesian optimization techniques. This scheme differs from *MethylNet*, as both the neural network topology and set of hyperparameters can be optimized through the application of successive Gaussian processes to update some prior of losses over the set of hyperparameters. However, the results presented in this manuscript utilized the randomized search design. The jobs can be launched in parallel and scaled to meet the demands of a larger compute cluster.

### Capsule Generation

Capsules specify the groupings of CpGs of the *MethylationArray* object. Capsule selection has been incorporated into the *hyperparameter_scan* and the *methylcaps-model* subcommands. Application Programming Interface (API) access to capsule selection and the building may be accessed through the *build_capsules* script. As mentioned in the results section, prespecified capsules include the following Illumina methylation array annotations – UCSC_RefGene_Name, UCSC_RefGene_Accession, UCSC_RefGene_Group, UCSC_CpG_Islands_Name, Relation_to_UCSC_CpG_Island, Phantom, DMR, Enhancer, HMM_Island, Regulatory_Feature_Name, Regulatory_Feature_Group, and DHS. Additionally, the following GSEA gene sets may be queried: C5.BP, C6, C1, H, C3.MIR, C2.CGP, C4.CM, C5.CC, C3.TFT, C5.MF, C7, C2.CP, C4.CGN. Users can also specify their own capsules through the presentation of a pickled dictionary containing a DataFrame that maps each CpG to a context name of choice. Capsule generation may also be accomplished by breaking up the entire hg19 genome into overlapping windows of fixed width [57] (*genomic_binned* selection). We recommend the utilization of the Circos tool [111] for visualization of derived capsule relationships using the *genomic_binned* option.

### Selection of Capsules for *MethylCapsNet* and *MethylSPWNet*

For the training of *MethylSPWNet*, we utilized all genes that overlapped with the 200,000 most variable CpGs across the CNS tumors. For *MethylCapsNet*, we could not utilize the complete set of genes due to the number of free parameters, a gene list of 650 genes was manually curated for *MethylCapsNet* that included genes related to WNT, SHH, DKK1, beta-catenin, SFRP, NPR3, amongst others. This list was reduced to 55 genes via recognition of genes by domain experts and thresholding of the minimum number of CpGs. As a further description, for both approaches, hyperparameter scans were utilized for pruning genes that did not contain a minimum number of CpGs (this threshold was varied via the hyperparameter scan), resulting in a lower number of genes than originally specified (n=10,341). In future iterations of the capsule-inspired network-based approach, gene selection constraints will be lifted via reduction of free parameters and the adoption of explicit network building approaches.

### Co-Methylation Embedding Modules

*MethylSPWNet* derived gene-level methylation summaries/embeddings were correlated to each other and within their own set of top genes. Louvain modularity was performed on a k-nearest neighbors graph of *MethylSPWNet* gene-level embeddings to establish preliminary modules. For *MethylSPWNet*, the final gene co-methylation/embedding analysis was carried out on a subtype-specific basis, done so through the use of WGCNA.

For *MethylCapsNet*, capsule-level embeddings were averaged across all individuals to form overall embeddings. Though just as relevant, these approaches can be extended to capsule-level embeddings on the individual level or aggregated across meaningful subgroups. To derive final measures of coordinated response between capsules, we averaged the routing matrix coefficients across the individuals to form a weighted bipartite graph and calculated a bipartite projection of the graph to form a unipartite graph of capsules. We utilized the Louvain modularity algorithm to discover hubs in this network and performed enrichment analyses on the pathway level using enrichr [112] to describe these hubs.

### Random Forest Approaches in Comparison

As a comparison to *MethylSPWNet* and *MethylCapsNet*, we adapted the Random Forest scheme featured in a DNAm machine learning classification study. We selected 10k CpGs by first fitting 100 random forest models, each themselves fit on 10k randomly selected CpGs. Shapley Additive Feature Explanations (SHAP) [113], was employed to determine the top CpGs from each random forest run. The 10k CpGs with the highest average rank across the 100 random forest models were selected for the final RF model. We note that we did not have access to the original set of 10k CpGs featured in the previous classifier development study. The previous study also utilized probability calibration methods to boost the model sensitivity and specificity, which we avoided to ensure a proper comparison between methods.

### Analysis of CpG Weights derived from *MethylSPWNet*

*MethylSPWNet* derives CpG-specific weights, 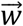, that relates each CpG to its respective gene. We rank-ordered, reverse rank-ordered, and absolute-value reverse rank-ordered these lists to yield CpGs that were important to differentiate the tumor types. We subset the first 1000 CpGs, marked which genes they corresponded to, and tested for enrichment using enrichr in our preliminary weight analysis. Finally, weights were also rank-ordered and reverse-rank ordered to yield the set of top negative and positive weights, respectively, the CpGs correspondent to the top number of CpGs (selected to highlight tendencies of enrichment and depletion) were related to the various island and gene context.

### Method to Cluster Gene-Level Brain Cancer Embeddings by Samples

Recall that embeddings for individuals for *MethylSPWNet* were given in the following form (the design matrix is of dimensionality samples by genes):

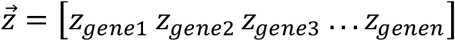

For *MethylNet*, the embeddings are derived using the encoder:

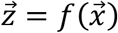

Embeddings for individuals using the *MethylCapsNet* approach (of dimensionality samples by genes by latent dimensions) can be obtained by either averaging or concatenating (an aggregation, or *AGG* operator) the gene-level embeddings:

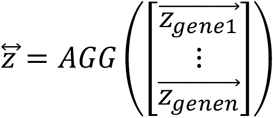

Stacking these vectors for individuals would yield a design matrix that can be clustered using methods such as hierarchical clustering. We implemented hierarchical clustering using *scikit-learn* (>0.22) and found 14 clusters to compare against true labels of cell-of-origin, histological subtype, and histological and molecular subtype using the v-measure statistic and cluster separation using the Silhouette coefficient.

### Web Application

We have developed a web application for the submission and investigation of *MethylCapsNet* outputs. The web application features three modules. The first is the network projection model, where capsules are related two to each other across subtypes, and network configurations can be changed by having some users tweak the relationships between the capsules and conservation. The second module displays routing information and the third module displays embedding information. Usage is detailed in the wiki.

### Analysis Hardware and Software

The analyses run for this work were optimized utilizing K80 GPUs at the Dartmouth Research Computing Cluster. The algorithms were designed using Python 3.7, PyTorch version 1.1, and CUDA 9.0.

### Dataset Preprocessing

We acquired data from GEO accession GSE109381 preprocessed data using *PyMethylProcess* and the subselected 200K of the most hypervariable CpGs (to focus on CpGs that may better differentiate CNS tumor subtypes) after functional normalization was applied to the data. SNPs and non-autosomal (sex chromosome) probes were omitted. Preprocessing steps have been detailed in using the pipeline of *PyMethylProcess* [21].

## Abbreviations

ANN: Artificial Neural Network
CGI: CpG Island
CIMP: CpG Island Methylator Phenotype
CNS: Central Nervous System
CpG: Cytosine Guanine Dinucleotide
DNAm: DNA Methylation
GBM: Glioblastoma Multiforme
LGG: Low-Grade Glioma
MB: Medulloblastoma
MLP: Multi-Layer Perceptron
ReLU: Rectified Linear Unit
Tanh: Hyperbolic Tangent
TDA: Topological Data Analysis
TOM: Topological Overlap Matrix
WGCNA: Weighted Gene Correlation Network Analysis

## Declarations

### Ethics Approval and Consent to Participate

Not applicable.

### Consent for Publication

Not applicable.

### Code Availability

The software (*MethylCapsNet, MethylSPWNet*) is open source and can be found on GitHub at https://github.com/Christensen-Lab-Dartmouth/MethylCapsNet, on PyPI under the tag *methylcapsnet*, and on Docker at *joshualevy44/methylcapsnet*. While new features may be developed for the *MethylCapsNet* framework, community contributions are welcome in the form of GitHub pull requests and issues. A test pipeline is available in the software implementation, a Wiki, help documentation, and example R scripts for possible downstream analyses can be found in the GitHub repository.

## Data Availability

Data used in this study was acquired from GEO accessions GSE109381, GSE84207, and GSE75067. Test data is available in our GitHub repository.

## Authors’ Contributions

The conception and design of the study were contributed by JJL and BCC. Implementation, programming, data acquisition, and analyses were by JJL. All authors contributed towards refining the analytic plan and direction. All authors contributed to the writing and editing of the manuscript. CLP and JJL tested the pipeline.

## Acknowledgments

We would like to acknowledge Christian Haudenschild, Hildreth Robert Frost, and A. James O’Malley for their thoughtful discussions.

