## Supplementary Materials for "MethylSPWNet and MethylCapsNet: Biologically Motivated Organization of DNAm Neural Network, Inspired by Capsule Networks"

**Supplementary Material**

**Gene-Level CNS Tumor Embeddings**

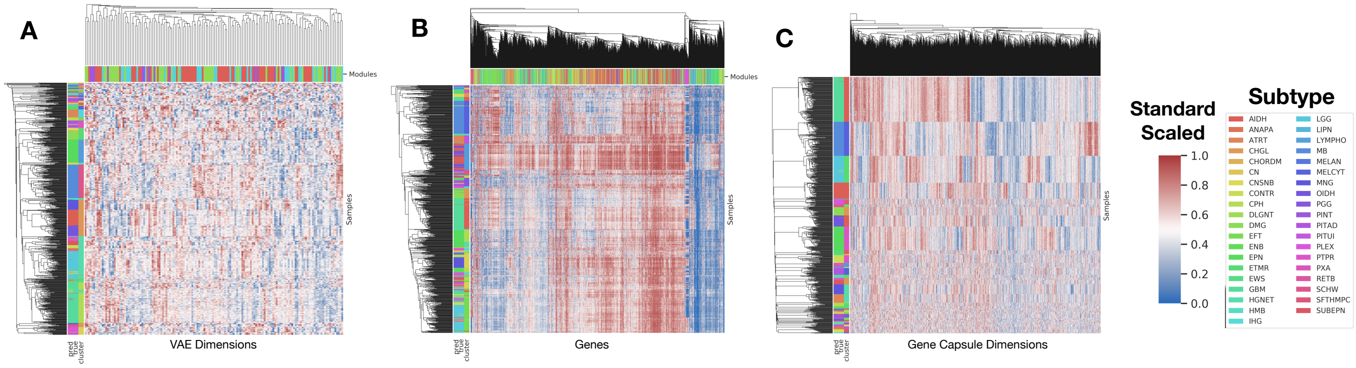


**Supplementary Figure 1**: Cluster heatmap plots of up to 2000 top components/capsules for: a) uninterpretable MethylNet dimensions, b) MethylSPWNet neural network defined gene methylation, c) 2200 dimensions comprising 55 MethylCapsNet capsules (55 capsules x 40 dimensions per Capsule = 2200 dimensions); samples constitute rows; capsules or latent dimensions constitute columns; rows are color labeled by predicted and true histological subtype and hierarchical clustering information; columns are color labeled by Louvain module assignment; all plots have been standard scaled to highlight patterns

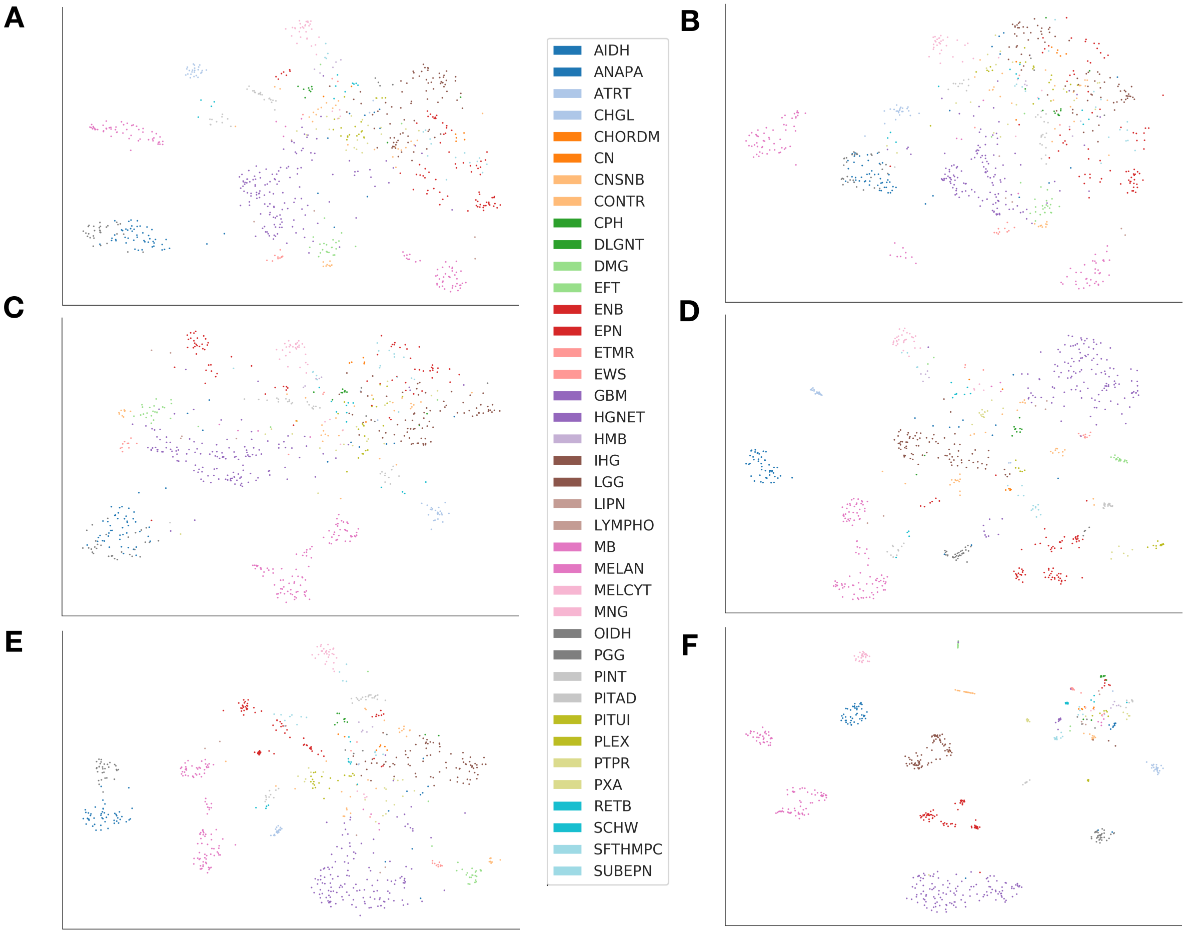

**Supplementary Figure 2:** NCVis Embeddings for a) all DNAm, b) Median gene DNAm, c) Median DNAm by gene context (upstream/downstream), d) *MethylNet*, e) *MethylSPWNet*, f) *MethylCapsNet*

**Supplementary Table 1:** Per-Subtype Breakdown of Classification Results for MethylSPWNet and MethylCapsNet on Test Set

| | MethylSPWNet | | | MethylCapsNet | | | |
|---|---|---|---|---|---|---|---|
| **Subtype** | **Precision** | **Recall** | **F1-Score** | **Precision** | **Recall** | **F1-Score** | **Support** |
| **AIDH** | 1.00 | 0.96 | 0.98 | 1.00 | 0.96 | 0.98 | 49 |
| **ANAPA** | 1.00 | 0.89 | 0.94 | 1.00 | 0.89 | 0.94 | 9 |
| **ATRT** | 1.00 | 1.00 | 1.00 | 1.00 | 1.00 | 1.00 | 24 |
| **CHGL** | 1.00 | 1.00 | 1.00 | 1.00 | 1.00 | 1.00 | 2 |
| **CHORDM** | 1.00 | 1.00 | 1.00 | 1.00 | 1.00 | 1.00 | 2 |
| **CN** | 1.00 | 1.00 | 1.00 | 1.00 | 1.00 | 1.00 | 5 |
| **CNSNB** | 1.00 | 1.00 | 1.00 | 1.00 | 1.00 | 1.00 | 9 |
| **CONTR** | 1.00 | 0.88 | 0.94 | 1.00 | 0.92 | 0.96 | 25 |
| **CPH** | 1.00 | 1.00 | 1.00 | 1.00 | 1.00 | 1.00 | 9 |
| **DLGNT** | 1.00 | 1.00 | 1.00 | 0.67 | 1.00 | 0.80 | 2 |
| **DMG** | 1.00 | 1.00 | 1.00 | 0.96 | 1.00 | 0.98 | 23 |
| **EFT** | 1.00 | 1.00 | 1.00 | 1.00 | 1.00 | 1.00 | 2 |
| **ENB** | 1.00 | 1.00 | 1.00 | 1.00 | 1.00 | 1.00 | 7 |
| **EPN** | 1.00 | 0.97 | 0.99 | 1.00 | 0.99 | 0.99 | 71 |
| **ETMR** | 0.91 | 1.00 | 0.95 | 0.91 | 1.00 | 0.95 | 10 |
| **EWS** | 1.00 | 1.00 | 1.00 | 1.00 | 1.00 | 1.00 | 3 |
| **GBM** | 0.98 | 1.00 | 0.99 | 0.98 | 0.99 | 0.98 | 130 |
| **HGNET** | 1.00 | 0.80 | 0.89 | 1.00 | 0.90 | 0.95 | 10 |
| **HMB** | 1.00 | 1.00 | 1.00 | 1.00 | 1.00 | 1.00 | 5 |
| **IHG** | 0.00 | 0.00 | 0.00 | 1.00 | 1.00 | 1.00 | 3 |
| **LGG** | 0.92 | 1.00 | 0.96 | 0.97 | 0.99 | 0.98 | 78 |
| **LIPN** | 1.00 | 1.00 | 1.00 | 1.00 | 1.00 | 1.00 | 2 |
| **LYMPHO** | 1.00 | 1.00 | 1.00 | 1.00 | 1.00 | 1.00 | 3 |
| **MB** | 1.00 | 1.00 | 1.00 | 1.00 | 1.00 | 1.00 | 100 |
| **MELAN** | 1.00 | 1.00 | 1.00 | 1.00 | 1.00 | 1.00 | 4 |
| **MELCYT** | 1.00 | 1.00 | 1.00 | 1.00 | 1.00 | 1.00 | 4 |
| **MNG** | 0.97 | 1.00 | 0.98 | 1.00 | 1.00 | 1.00 | 29 |
| **OIDH** | 0.94 | 1.00 | 0.97 | 0.94 | 1.00 | 0.97 | 31 |
| **PGG** | 1.00 | 1.00 | 1.00 | 1.00 | 1.00 | 1.00 | 4 |
| **PINT** | 1.00 | 0.90 | 0.95 | 0.91 | 1.00 | 0.95 | 10 |
| **PITAD** | 1.00 | 1.00 | 1.00 | 1.00 | 1.00 | 1.00 | 20 |
| **PITUI** | 1.00 | 1.00 | 1.00 | 1.00 | 1.00 | 1.00 | 6 |
| **PLEX** | 1.00 | 1.00 | 1.00 | 1.00 | 1.00 | 1.00 | 15 |

| | | | | | | | |
|---|---|---|---|---|---|---|---|
| **PTPR** | 1.00 | 1.00 | 1.00 | 1.00 | 1.00 | 1.00 | 6 |
| **PXA** | 1.00 | 0.92 | 0.96 | 0.92 | 0.85 | 0.88 | 13 |
| **RETB** | 0.80 | 1.00 | 0.89 | 1.00 | 0.75 | 0.86 | 4 |
| **SCHW** | 1.00 | 1.00 | 1.00 | 1.00 | 1.00 | 1.00 | 8 |
| **SFTHMPC** | 1.00 | 1.00 | 1.00 | 1.00 | 1.00 | 1.00 | 4 |
| **SUBEPN** | 1.00 | 1.00 | 1.00 | 1.00 | 1.00 | 1.00 | 13 |
| **Overall Accuracy** | | | 0.98 | | | 0.98 | |
| **Macro-Average** | 0.96 | 0.96 | 0.96 | 0.98 | 0.98 | 0.98 | 754 |
| **Weighted Average** | 0.98 | 0.98 | 0.98 | 0.98 | 0.98 | 0.98 | 754 |

**Supplementary Table 2:** Concordance Statistic from Cluster Assignment (k=14) to: cell of origin, molecular subspecification, and histology

| Method | Cell of Origin | Histology+Molecular | Histology |
|---|---|---|---|
| **MethylNet** | 0.72±0.012 | 0.68±0.0069 | 0.8±0.0097 |
| **MethylSPWNet** | 0.73±0.011 | 0.72±0.0059 | 0.81±0.0086 |
| **MethylCapsNet** | 0.65±0.012 | 0.66±0.0068 | 0.8±0.0078 |
| **Median Methylation** | 0.11±0.014 | 0.15±0.016 | 0.17±0.019 |
| **Median Methylation by Context** | 0.11±0.014 | 0.15±0.016 | 0.17±0.019 |

### Preliminary Pathways and Module Analysis

To identify modules of gene co-methylation patterns and understand how they relate to underlying pathways we selected the 2000 most variably methylated genes across the 38 brain cancer subtypes as defined by gene median methylation and SPW derived gene-level methylation. We clustered the resulting genes using Louvain Modularity to identify co-methylation modules and then tested for enrichment after combining the two largest modules. The genes that were identified in the largest 2 modules were selected for enrichment analysis. We also performed a WGCNA analysis on the genes to identify key modules, the results of which have been included in the main text. Results indicate that these top two modules are highly related to key cancer signaling pathways, including Rap1, Ras, WNT, and GABAergic pathways, to name a few. Alternative specification of key genes using the same module detection analysis for median methylation yielded no related pathways, whereas ranking genes by median absolute deviation recovered many pathways identified by *MethylSPWNet*, though not at the same power as that derived from the neural network. Furthermore, since the neural network pathway enrichment analysis discovered pathways with higher power than that specified using median methylation, it detected pathways with less well-known associations, such as Type I Diabetes Mellitus (hsa04940), corroborated by the literature [1]. Additionally, Type 1 diabetes is an autoimmune disease; we found associations with several immune evading markers, which may account for the spurious associations. All pathway enrichment tests are available in Additional File 4.

Then, we rank-ordered genes by their maximum weight from the largest weight to the smallest and rank-ordered the minimum weights from smallest to largest, extracting the first thousand CpGs from each list and their associated genes. As such, two separate enrichment analyses were conducted, one for the negative weights and one for the positive weights. Many similar, but not intersecting pathways were found to be statistically important for each of these analyses, as compared to the modularity-based analyses. Particularly, some new pathways include axon guidance, Hippo signaling, Circadian entrainment, melanogenesis, TGF-B, and MAPK pathways. Complete results can be found in the additional spreadsheet. An analysis of the top 1000 hypervariable methylated CpGs across the cancer type yielded pathways that overlapped with *MethylSPWNet* pathways, even though median absolute variation was not associated with the weight assigned to individual CpGs (Supplementary Figure 5; Additional File 4).

We found that positive CpG weights tended to be associated with islands, North Shores, 5'UTR, and TSS200, while negative neural network derived CpG weights are associated with gene body, 3'UTR and possibly South shelves. Overall, there were associations with islands, 5'UTR, and TSS200. Aggregating by context into upstream sites (5'UTR, TSS200, TSS1500) and downstream sites (1$^{st}$ Exon, Gene Body, 3'UTR), upstream sites appear to be associated with positive neural network weights, while downstream overall appear to be associated with negative neural network weights.

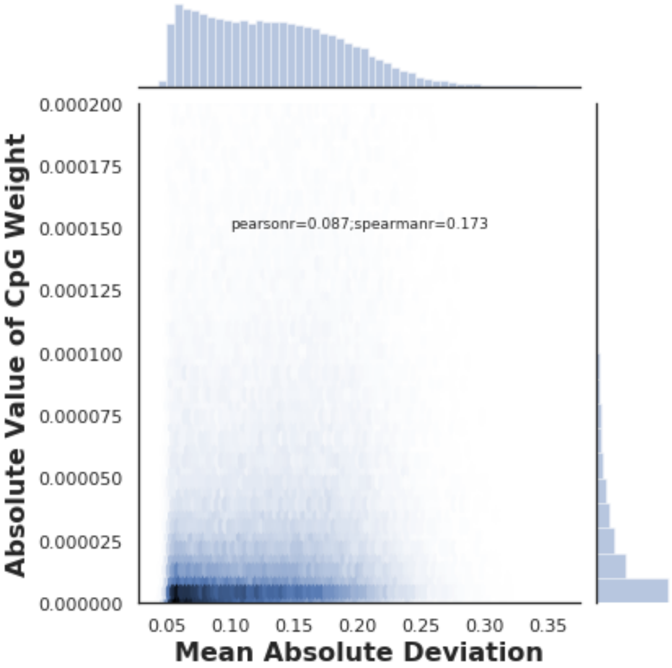


**Supplementary Figure 3:** Relationship of mean absolute deviation in DNAm across samples and CpG weight

**Supplementary Table 3:** Summarized results from preliminary pathway analyses (See additional file 4)

| Summary of Notable KEGG Pathways | Notable Genes (n=9 sampled) | Average OncoScore | Gene Set Score |
|---|---|---|---|
| **Ras signaling pathway Homo sapiens hsa04014** | ABL1;PDGFRA;EGFR;EPHA2;ABL2;PRKACA;RALB;ETS2;PAK6 | 87.07 | 21.79 |

| | | | |
|---|---|---|---|
| **Pathways in cancer Homo sapiens hsa05200** | ABL1;CCND1;ZBTB16;MECOM;PDGFRA;CDK6;PTEN;RUNX1;CCNA1 | 87.07 | 18.80 |
| **Hippo signaling pathway Homo sapiens hsa04390** | LATS2;TP73;LIMD1;WWTR1;CTNNA1;LLGL2;DLG2;WNT5A;PARD3 | 82.93 | 31.10 |
| **Axon guidance Homo sapiens hsa04360** | NTN1;EPHA2;UNC5A;EFNA1;PAK6;EPHB4;EPHA3;EPHB3;SLIT3 | 82.89 | 19.89 |
| **Phospholipase D signaling pathway Homo sapiens hsa04072** | TSC2;EGFR;RALB;RHEB;MAPK1;GAB2;RALGDS;GNAS;PLD1 | 79.28 | 16.64 |
| **Focal adhesion Homo sapiens hsa04510** | CCND1;PTEN;EGFR;ITGB4;MAPK1;VAV2;ITGB5;HGF;RAPGEF1 | 78.17 | 13.30 |
| **MAPK signaling pathway Homo sapiens hsa04010** | MECOM;PDGFRA;EGFR;DAXX;MAP2K2;RPS6KA2;FGFR3;PRKCA;PDGFRB | 76.62 | 20.23 |
| **Gap junction Homo sapiens hsa04540** | PDGFRA;EGFR;MAP2K2;PRKCA;PDGFRB;PRKCB;GNAS;PDGFB;ADCY9 | 76.33 | 33.17 |
| **Prostate cancer Homo sapiens hsa05215** | PDGFRA;EGFR;MAP2K2;CREB5;PDGFRB;CREBBP;TCF7L1;TCF7L2;PDGFB | 76.33 | 27.64 |
| **Melanoma Homo sapiens hsa05218** | PDGFRA;CDK6;EGFR;MITF;PDGFRB;FGF2;FGF1 | 76.33 | 26.57 |
| **Calcium signaling pathway Homo sapiens hsa04020** | PDGFRA;EGFR;STIM1;PRKCA;PDGFRB;ITPKA;PRKCB;GNAS;CACNA1G | 76.33 | 22.42 |
| **Rap1 signaling pathway Homo sapiens hsa04015** | PDGFRA;EGFR;EPHA2;EFNA1;SIPA1;RALB;MAP2K2;BCAR1;MAPK1 | 76.33 | 20.27 |
| **Type I diabetes mellitus Homo sapiens hsa04940** | TNF;PRF1;CD28;IL1A;LTA;ICA1;HLA-DRB5;HLA-DMA;HLA-C | 73.23 | 30.35 |
| **Adherens junction Homo sapiens hsa04520** | EGFR;CTNNA1;PTPN6;CREBBP;PTPRF;PARD3;TCF7L1;SMAD3;TCF7L2 | 69.06 | 69.62 |
| **Estrogen signaling pathway Homo sapiens hsa04915** | EGFR;MAP2K2;CREB5;GNAS;HSPA1L;ADCY9;ADCY4;FKBP5;GABBR1 | 69.06 | 24.66 |
| **Vascular smooth muscle contraction Homo sapiens hsa04270** | MYH11;MAP2K2;PRKCB;GNAS;ADCY9;ADCY4;PPP1R14A;PRKG1;ADRA1D | 66.53 | 22.45 |
| **Arrhythmogenic right ventricular cardiomyopathy (ARVC) Homo sapiens hsa05412** | CTNNA1;ITGB4;ITGA11;TCF7L1;ITGB7;TCF7L2;CTNNA2;ITGA8;SLC8A1 | 64.02 | 26.65 |
| **Cell adhesion molecules (CAMs) Homo sapiens hsa04514** | CADM1;NCAM1;CDH3;PTPRF;CLDN9;CDH4;ITGB7;CADM3;ITGA8 | 60.56 | 17.10 |
| **Glutamatergic synapse Homo sapiens hsa04724** | GLS2;PRKACA;MAPK1;PRKCA;PRKCB;GNAS;GNG7;GNG12;PLD1 | 56.32 | 22.85 |
| **GABAergic synapse Homo sapiens hsa04727** | GLS2;PRKACA;GNG7;GNG12;GABRP;GNB1;GNG3;PRKCG;ADCY5 | 56.32 | 19.37 |
| **Serotonergic synapse Homo sapiens hsa04726** | PRKACA;MAPK1;GNAS;GNG7;GNG12;GNB1;ALOX12B;PLA2G4B;GNG3 | 55.18 | 25.83 |
| **Morphine addiction Homo sapiens hsa05032** | PRKACA;GNAS;GNG7;GNG12;GABRP;GNB1;GNG3;PRKCG;OPRM1 | 55.18 | 17.89 |
| **Retrograde endocannabinoid signaling Homo sapiens hsa04723** | PRKACA;MAPK1;GNG7;GNG12;GABRP;GNB1;GNG3;PRKCG;ADCY5 | 55.18 | 17.66 |
| **Melanogenesis Homo sapiens hsa04916** | MAP2K2;MITF;PRKCA;PRKCB;GNAS;CREBBP;TCF7L1;FZD6;TCF7L2 | 52.87 | 30.49 |
| **Wnt signaling pathway Homo sapiens hsa04310** | CACYBP;PRKCA;PRKCB;CREBBP;TCF7L1;FZD6;SMAD3;TCF7L2;PRICKLE2 | 50.38 | 23.72 |
| **Dilated cardiomyopathy Homo sapiens hsa05414** | ITGB4;GNAS;ITGA11;ITGB7;TGFB2;ADCY2;ADCY9;ADCY8;ADCY4 | 50.36 | 20.14 |
| **Aldosterone synthesis and secretion Homo sapiens hsa04925** | CREB5;PRKCA;PRKCB;GNAS;CACNA1G;ADCY2;NR4A2;CAMK1D;CALML5 | 48.99 | 34.21 |
| **Circadian entrainment Homo sapiens hsa04713** | PRKCA;PRKCB;GNAS;CACNA1G;ADCY2;GNB1;CALML5;ADCY9;ADCY8 | 46.73 | 29.09 |
| **TGF-beta signaling pathway Homo sapiens hsa04350** | CREBBP;TGIF1;ACVR1C;SMAD3;SMAD7;TGFB2;INHBB;BMP7;PITX2 | 39.35 | 26.16 |

### Context Enrichment Tables and Figures

We sought to evaluate genomic contexts independently that bring about patient phenotype. CpGs with the most positive neural network weights were found to be enriched for CpG islands (OR = 1.53; p-value<0.01), south shelves (OR=1.52; p=0.04), open sea regions (OR=1.18; p<0.01), 5'UTR regions (OR=1.52; p=0.05), and the first exon (OR=1.62; p=0.02). Positive weight CpGs were found to be depleted in north shores (OR=0.54; p<0.01), south shores (OR=0.63; p=0.05), gene bodies (OR=0.75; p=0.05) and TSS1500 (OR=0.72; p=0.02). Negative neural network weight CpGs were found to be enriched in north shelves (OR=2.47; p=0.05), north shores (OR=1.23; p=0.01), TSS200 (OR=1.24; p=0.05) and 5'UTR (OR=1.26; p=0.01). These negative weight CpGs were depleted in south shelf regions (OR=0.49; p=0.02), south shores (OR=0.81; p=0.04) and gene bodies (OR=0.86; p=0.02).

**Supplementary Table 4:** Fischer’s Exact tests for enrichment of positive neural network weights for particular contexts

| | Number of Highest Positive Weighted CpGs | | | | | |
|---|---|---|---|---|---|---|
| | 250 | | 500 | | 2100 | |
| | Odds Ratio | P-Value | Odds Ratio | P-Value | Odds Ratio | P-Value |
| **Island** | **1.53** | **0.00** | 1.05 | 0.64 | 0.95 | 0.29 |
| **N_Shore** | **0.54** | **0.00** | 0.96 | 0.80 | 0.94 | 0.38 |
| **N_Shelf** | 1.49 | 0.11 | 1.03 | 0.84 | 0.85 | 0.14 |
| **S_Shore** | **0.63** | **0.05** | 0.76 | 0.08 | 0.92 | 0.26 |
| **S_Shelf** | 1.23 | 0.42 | **1.52** | **0.04** | 0.86 | 0.24 |
| **OpenSea** | 0.93 | 0.60 | 1.00 | 1.00 | **1.18** | **0.00** |
| | **Number of Highest Negative Weighted CpGs** | | | | | |
| | **50** | | **500** | | **1200** | |
| | Odds Ratio | P-Value | Odds Ratio | P-Value | Odds Ratio | P-Value |
| **Island** | 0.72 | 0.42 | 1.14 | 0.20 | 0.99 | 0.95 |
| **N_Shore** | 1.18 | 0.70 | 0.93 | 0.62 | **1.23** | **0.01** |
| **N_Shelf** | **2.47** | **0.05** | 0.91 | 0.76 | 0.81 | 0.16 |
| **S_Shore** | 0.90 | 1.00 | 1.04 | 0.77 | **0.81** | **0.04** |
| **S_Shelf** | 2.13 | 0.13 | **0.49** | **0.02** | **0.68** | **0.03** |
| **OpenSea** | 0.76 | 0.39 | 1.02 | 0.85 | 1.05 | 0.38 |
| | **Number of Highest Positive Weighted CpGs** | | | | | |
| | **150** | | **200** | | **500** | |
| | Odds Ratio | P-Value | Odds Ratio | P-Value | Odds Ratio | P-Value |
| **TSS1500** | 0.64 | 0.09 | 0.77 | 0.24 | **0.72** | **0.02** |
| **TSS200** | 1.21 | 0.46 | 1.21 | 0.44 | 1.07 | 0.63 |
| **5'UTR** | **1.52** | **0.05** | **1.42** | **0.05** | 1.07 | 0.61 |
| **1stExon** | 1.46 | 0.11 | **1.62** | **0.02** | 1.20 | 0.19 |
| **Body** | 0.79 | 0.16 | **0.75** | **0.05** | 1.03 | 0.75 |
| **3'UTR** | 0.85 | 0.85 | 0.63 | 0.31 | 0.98 | 1.00 |

| | Number of Highest Negative Weighted CpGs | |
|---|---|---|
| | **1000** | |
| | Odds Ratio | P-Value |
| **TSS1500** | 0.93 | 0.45 |
| **TSS200** | **1.24** | **0.05** |
| **5'UTR** | **1.26** | **0.01** |
| **1stExon** | 0.91 | 0.40 |
| **Body** | **0.86** | **0.02** |
| **3'UTR** | 1.17 | 0.29 |

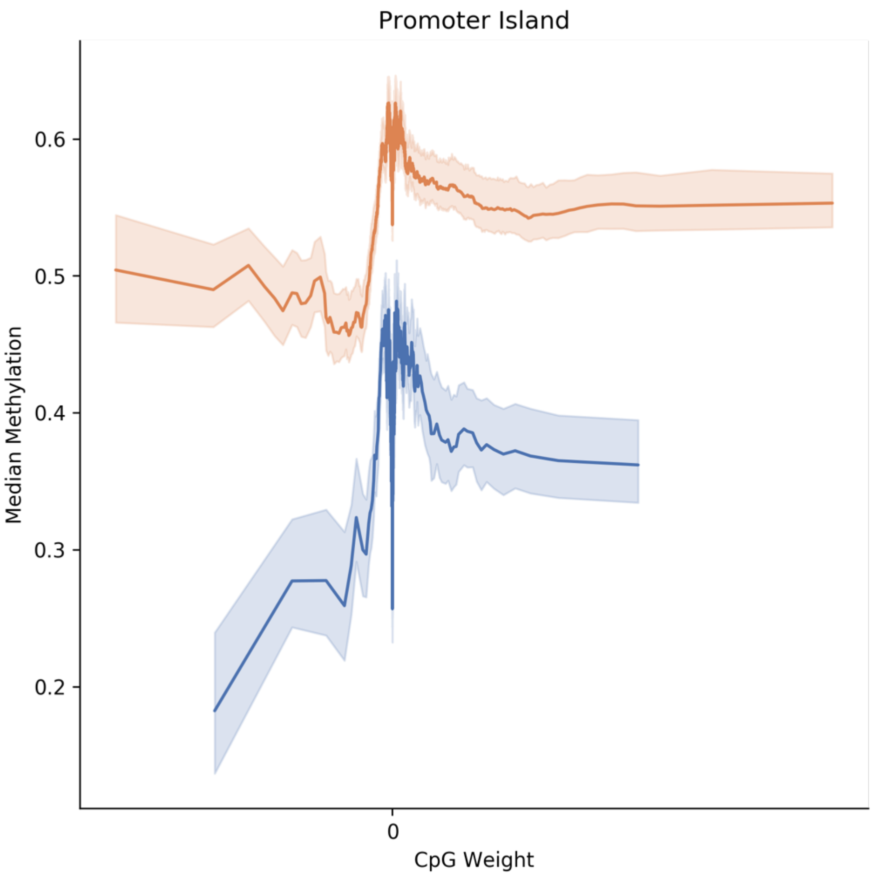


**Supplementary Figure 4:** Median methylation vs CpG weight; stratified by CpGs associated with island promoter (orange) and sites that are not associated with island promoters (blue); non-parametric bootstrapping was conducted across median aggregates for each tumor subtype to derive confidence intervals; note strong dip in median methylation near CpG weight of 0

**Screenshot of Web Application and Network Plots**

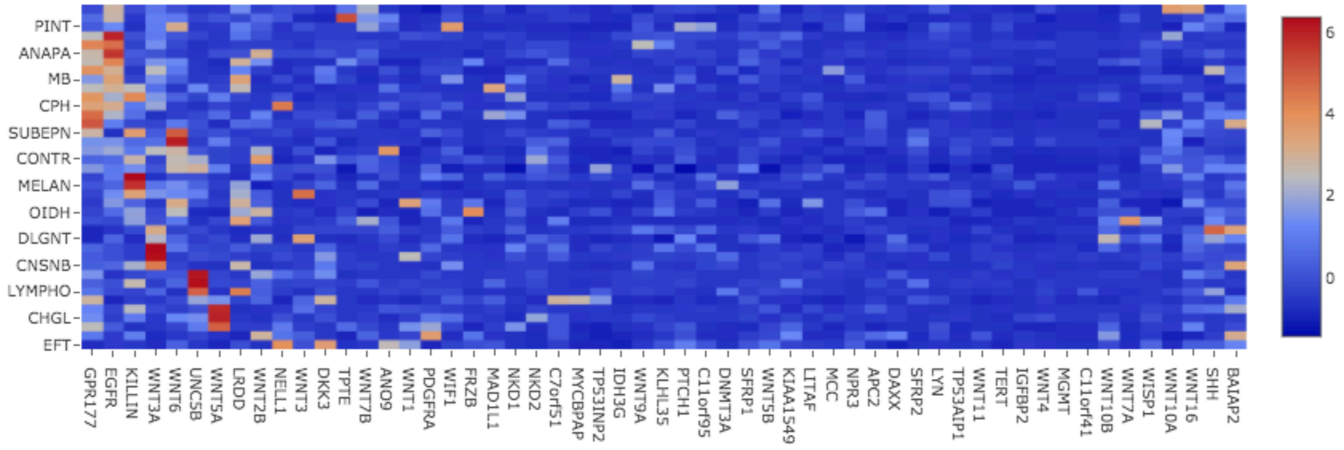


**Supplementary Figure 5:** Bipartite routing matrix that relates primary gene capsules to disease subtypes; red values indicate strong positive associations

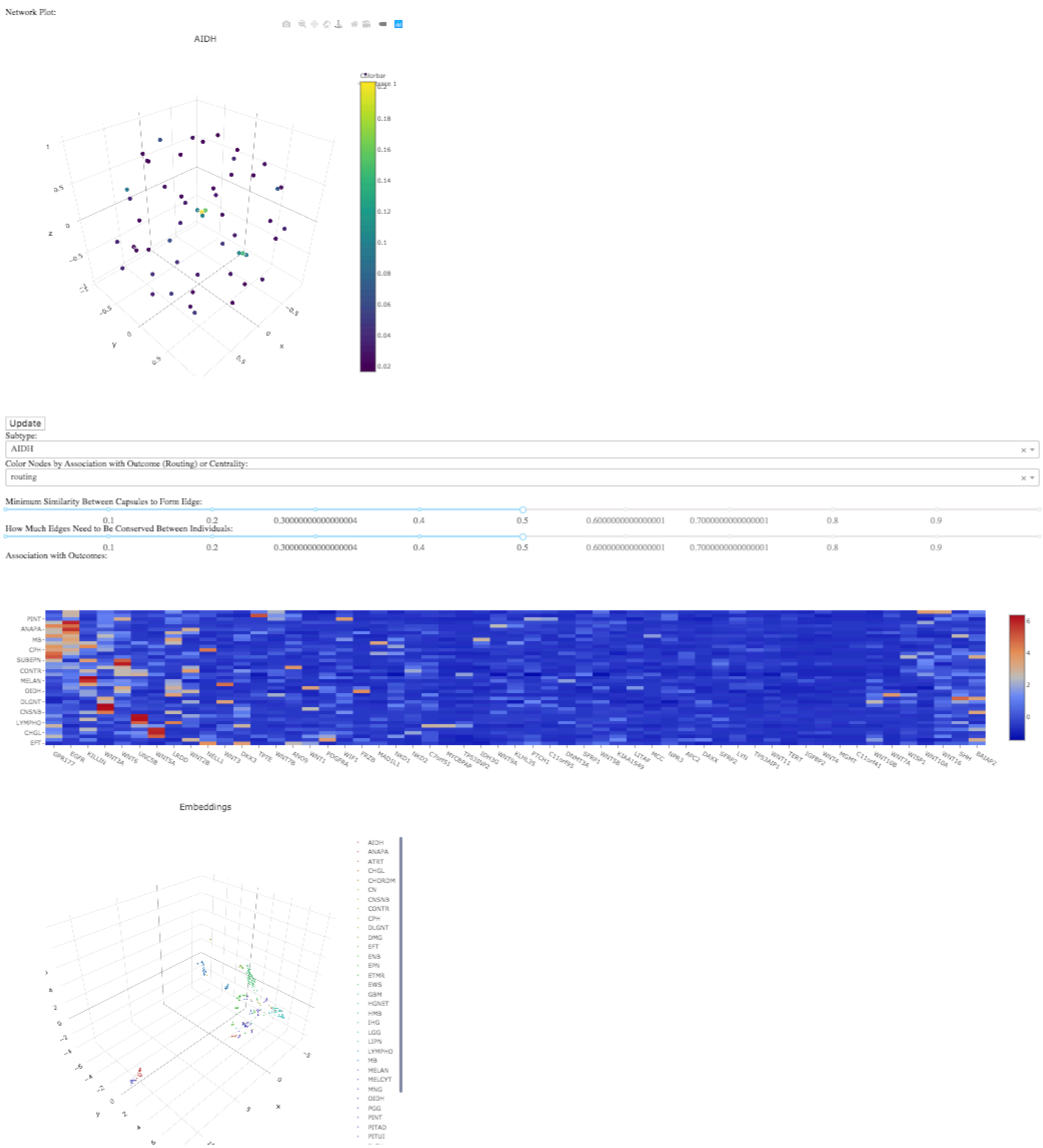


**Supplementary Figure 6:** Screenshot of entire web application interface where results are uploaded to; comprised of network discovery, routing matrix and embedding profile components

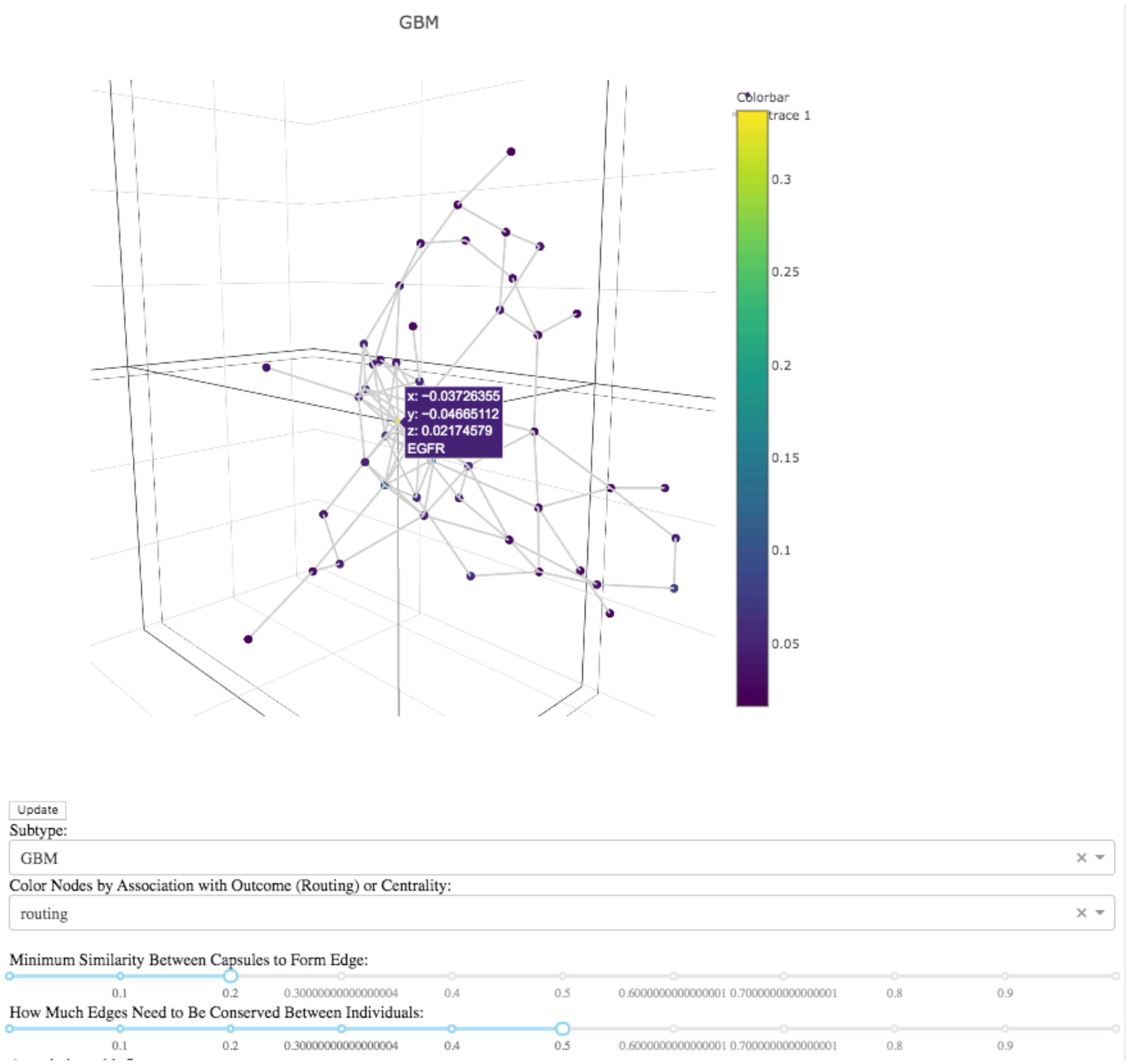


**Supplementary Figure 7:** Web application forms network using MethylCapsNet for GBM and uncovers EGFR as the most related gene using via the gene's routing coefficient

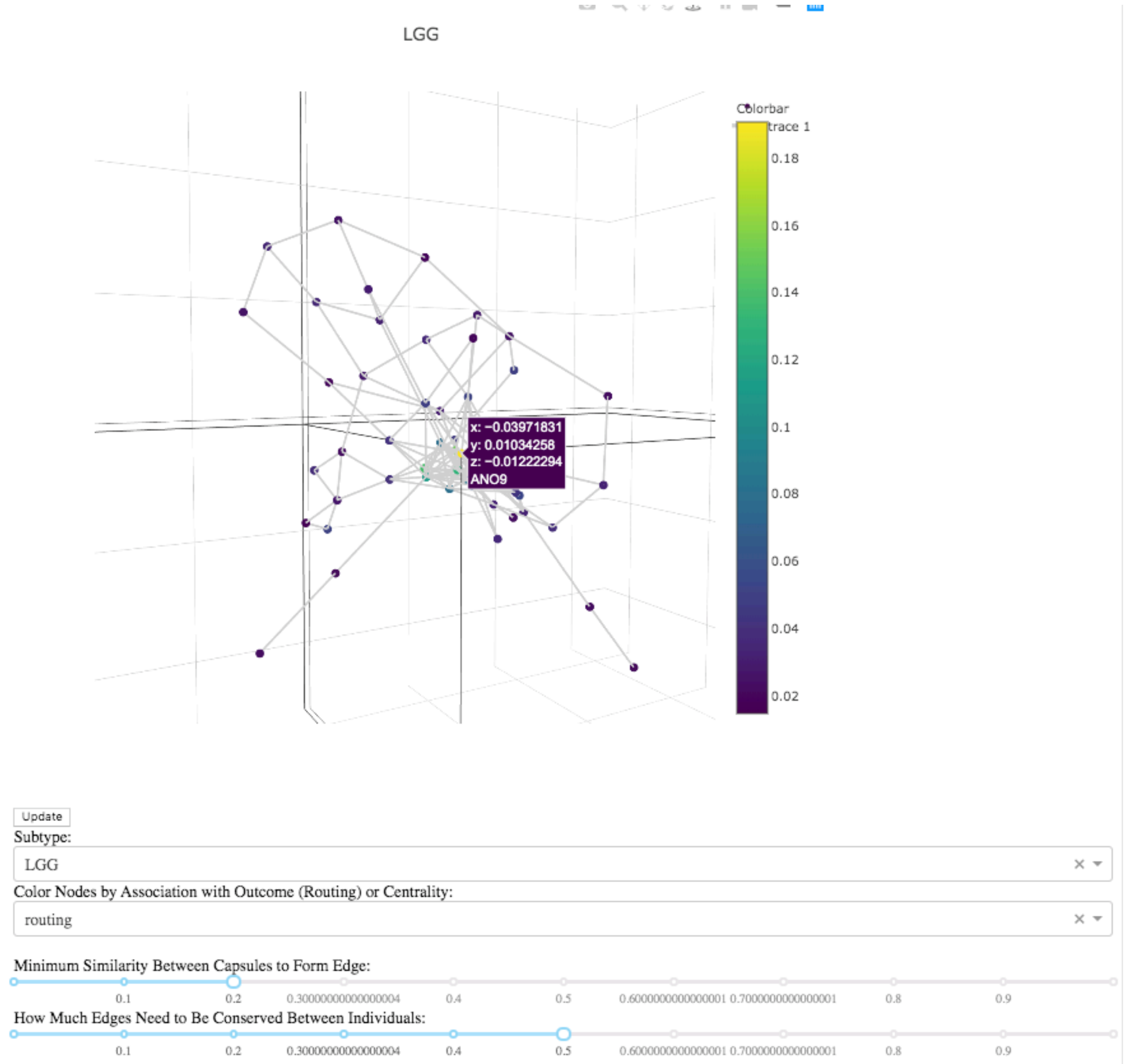


**Supplementary Figure 8:** Web application forms network using MethylCapsNet for LGG and uncovers ANO9 as the most related gene using via the gene's routing coefficient; previously implicated in [2] and related to the EGFR pathway

### Alternative Capsule Formations and Cancer Subtypes

Here, we report classification results for an alternative set of datasets and capsules, as detailed in the results section: "Exploration of Alternative Capsule Formations and Cancer Subtypes". *Half Genes* are intersected with the following set of genes: https://github.com/Christensen-Lab-Dartmouth/MethylCapsNet/blob/master/test_data/brain_cancer_gene_list.txt . *New Genes* does not intersect with any of these genes. *Curated Genes* intersects with the following set of genes: https://github.com/Christensen-Lab-Dartmouth/MethylCapsNet/blob/master/test_data/breast_cancer_gene_list.txt

**Supplementary Table 5:** Classification performance for alternative selection of capsules; classification performance is macro-averaged and 95% confidence intervals estimated using 1000-sample non-parametric bootstrap

| Dataset | Capsule Type | Algorithm | Accuracy | Precision | Recall | F1-Score |
|---|---|---|---|---|---|---|
| **CNS Tumor** | Half Genes | MethylCapsNet | 0.98±0.0045 | 0.98±0.01 | 0.98±0.011 | 0.98±0.011 |
| | New Genes | MethylCapsNet | 0.98±0.0048 | 0.98±0.013 | 0.97±0.013 | 0.97±0.013 |
| | Genomic Binned | MethylCapsNet | 0.99±0.0043 | 0.99±0.01 | 0.97±0.012 | 0.98±0.011 |
| **Breast Cancer** | Curated Genes | MethylCapsNet | 0.79±0.029 | 0.84±0.035 | 0.71±0.035 | 0.75±0.036 |
| | Genomic Binned | MethylCapsNet | 0.75±0.031 | 0.74±0.035 | 0.75±0.036 | 0.74±0.034 |
| | Island Promoters | MethylSPWNet | 0.75±0.031 | 0.78±0.039 | 0.7±0.035 | 0.72±0.037 |
| | N/A | MethylNet | 0.77±0.03 | 0.75±0.033 | 0.78±0.033 | 0.76±0.032 |

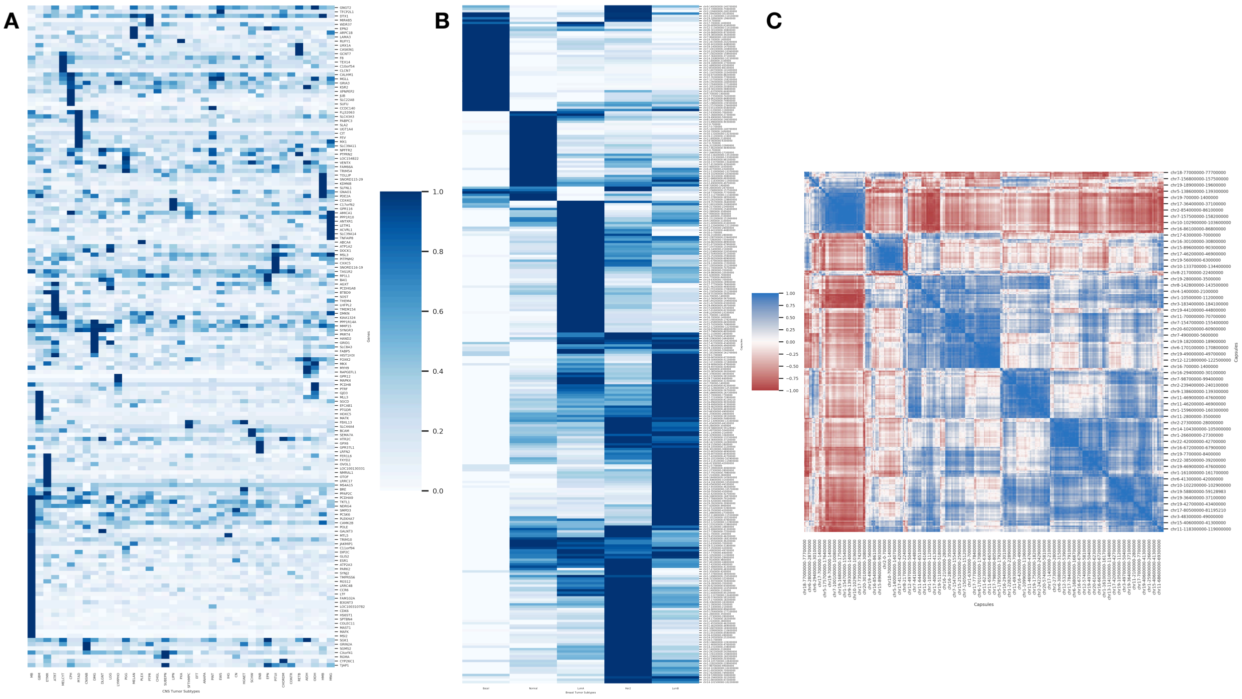


**Supplementary Figure 9:** Heatmaps of routing matrices (a,b; rows are capsules, columns are genes) and unipartite projection of routing matrix (c; rows and columns are capsules) for the following scenarios: a) *Half Genes* for CNS tumor classification, b-c) 700kb non-overlapping *Genomic Regions* for PAM50 molecular classification

**Supplementary Table 6:** Set of genes utilized by MethylCapsNet for CNS classification; first set of genes reference set used for original classification task in paper; second two sets reference alternative sets of capsules that were formed; set of genes were selected from larger list of genes, but only included if they each have at least 10 annotated CpGs; gene list selection was done through inclusion/exclusion of a larger list of genes (eg. all/half/none found here: https://github.com/Christensen-Lab-Dartmouth/MethylCapsNet/blob/master/test_data/brain_cancer_gene_list.txt) and all genes

| **MethylCapsNet Original Genes** | | | | |
|---|---|---|---|---|
| IDH3G | WNT7A | EGFR | LRDD | WNT1 |
| WNT3A | WNT5A | SHH | NELL1 | WIF1 |
| GPR177 | PDGFRA | C7orf51 | ANO9 | NKD1 |
| WNT2B | SFRP2 | WNT16 | KLHL35 | LITAF |
| WNT4 | NPR3 | SFRP1 | DKK3 | BAIAP2 |
| WNT9A | TERT | WISP1 | TP53AIP1 | WNT3 |
| WNT6 | MCC | LYN | C11orf95 | MYCBPAP |
| DNMT3A | NKD2 | PTCH1 | WNT11 | APC2 |
| WNT10A | DAXX | MGMT | C11orf41 | TP53INP2 |
| FRZB | KIAA1549 | KILLIN | WNT10B | TPTE |
| IGFBP2 | MAD1L1 | UNC5B | WNT5B | WNT7B |
| **MethylCapsNet Modified/Half Genes** | | | | |
| MAP3K15 | MEIS1 | KIAA1549 | DEAF1 | BAIAP2 |
| SYP | IGFBP2 | GRM8 | TP53AIP1 | MGAT5B |
| FAM122C | CTDSPL | CAMK2B | USP2 | TTYH2 |
| IDH3G | NISCH | C7orf51 | ELF5 | FAM171A2 |
| RNF207 | WNT7A | GNA12 | C11orf95 | RICH2 |
| DUSP27 | WNT5A | LOC642006 | WNT11 | MPO |
| FGR | ZNF827 | FASTK | C11orf92 | KCNAB3 |
| WNT3A | GPM6A | FOXP2 | HOXC5 | C17orf104 |
| GPR177 | NPFFR2 | SFRP1 | FBRSL1 | MIR33B |
| WNT2B | C4orf42 | KCNB2 | DTX1 | C17orf77 |
| LGALS8 | NPR3 | WISP1 | WNT5B | C19orf6 |
| ALPL | TERT | DOCK5 | WNT1 | DLL3 |
| C1orf61 | MCC | NDRG1 | CRIP2 | PDE4A |
| WNT6 | NKD2 | ZMAT4 | SLC7A8 | MYH14 |
| BIN1 | SHROOM1 | PAX5 | C14orf23 | HAS1 |
| UXS1 | LEMD2 | MGMT | CGNL1 | ZNF135 |
| ASB18 | PHACTR2 | C10orf108 | KIAA0182 | RELB |
| WNT10A | C6orf25 | UNC5B | NKD1 | MMP9 |
| NR4A2 | UBD | PPYR1 | SULT1A1 | ZNF334 |
| BOLL | BAT3 | BLNK | LITAF | SCUBE1 |
| STK25 | TNF | EFCAB4A | HSF4 | PIWIL3 |
| **MethylCapsNet Other/New Genes** | | | | |
| HTR2C | JAKMIP1 | CDK6 | C11orf94 | GLIS2 |
| TKTL1 | RGS12 | MAFK | AMICA1 | KDM6B |
| XPNPEP2 | HAND2 | FBXL13 | SLC22A8 | ATP2A3 |
| F8 | LETM1 | ARPC1B | CIT | RAPGEFL1 |
| GRIA3 | NPFFR2 | LRRC17 | HOXC5 | MSI2 |
| MSL3 | TMEM154 | MLL3 | KSR2 | C17orf62 |
| CXorf41 | PCDHGA8 | BAI1 | DTX1 | TEX14 |
| KIAA1324 | RUFY1 | RP1L1 | POLE | PTRF |
| SLFNL1 | PCDHA9 | SLC39A14 | PITPNM2 | GNGT2 |
| LMX1A | LHFPL2 | FER1L6 | ACVRL1 | GJD3 |
| TAS1R2 | LOC100310782 | FAM66A | PABPC3 | EPN2 |
| GPR37L1 | TNFAIP8 | FABP5 | PCDH8 | FOXK2 |
| ABCA4 | CXXC5 | EFCAB1 | GPR12 | SLC39A11 |
| ATP1A2 | SGCD | FAM102A | JUB | SOST |
| LOC100130331 | GPX6 | DOCK1 | SLC8A3 | MAPK4 |
| THEM4 | SLC44A4 | DIP2C | MIR485 | LAMA3 |
| OTOF | TJAP1 | VENTX | PTGDR | LRRC4B |
| FEV | PARK2 | CALHM1 | SNORD116-19 | SPTBN4 |
| BRE | PPP1R10 | WDR37 | SEMA7A | MAST1 |
| FLJ32063 | BTBD9 | CYP26C1 | SNORD115-29 | B3GNT3 |
| HS6ST1 | TRIM10 | GRID1 | RGMA | MATK |
| ANTXR1 | GPR116 | MKX | PCSK6 | PPAP2C |
| COLEC11 | SYNJ2 | SUFU | GRIN2A | BCAM |
| AGXT | ESR1 | C10orf54 | CLCN7 | PPP1R14A |
| UGT1A4 | SGK1 | PDE2A | CASKIN1 | DMKN |
| TFCP2L1 | LRFN2 | PLEKHA7 | MMP15 | SLA2 |
| CCDC140 | CCR6 | SLC43A3 | GNAO1 | GCNT7 |
| GALNT3 | HIST1H3I | MTL5 | NDRG4 | COX4I2 |
| TRIM54 | PTPRN2 | TOLLIP | NMRAL1 | MX1 |
| MGLL | CAMK2B | OVOL1 | SYNGR3 | TMPRSS6 |
| LTF | LOC154822 | FXYD2 | SMPD3 | MYH9 |
| SGMS2 | PRRT4 | MS4A15 | | |

### Description of Additional Files

Appended to the supplementary material are a collection of files which demonstrate what kind of results to expect if running a differential embedding analysis, followed by a pathway and WGCNA analysis. The first tab of files 1-3 indicate results from an empirical bayes analysis. The second tab demonstrates pathway analyses of results from the first tab. Finally, tab 3 displays results from the WGCNA analysis. The first row indicates the module name, while the second row indicates the correlation with the subtype. The third row displays the significance of such association and remaining rows display top 20 hub genes for the WGCNA modules.

The fourth additional file contains preliminary pathway results (see "Preliminary Pathways and Module Analysis"), where sets of genes were input into enrichr based on cutoff criteria discussed in the manuscript and supplementary material. The first tab displays pathways associated with genes that were produced by whether they contained some of the lowest CpG weights from MethylSPWNet. The second tab displays pathways for genes associated with the highest CpG weights from MethylSPWNet. The third tab displays pathways for genes that associated with hypervariablly methylated CpGs (as defined by median absolute deviation) across all cancer subtypes.
